# Virtual Genome Walking: Generating gene models for the salamander *Ambystoma mexicanum*

**DOI:** 10.1101/185157

**Authors:** Teri Evans, Andrew Johnson, Matt Loose

## Abstract

Large repeat rich genomes present challenges for assembly and identification of gene models with short read technologies. Here we present a method we call Virtual Genome Walking which uses an iterative assembly approach to first identify exons from *de-novo* assembled transcripts and assemble whole genome reads against each exon. This process is iterated allowing the extension of exons. These linked assemblies are refined to generate gene models including upstream and downstream genomic sequence as well as intronic sequence. We test this method using a 20X genomic read set for the axolotl, the genome of which is estimated to be 30 Gb in size. These reads were previously reported to be effectively impossible to assemble. Here we provide almost 1 Gb of assembled sequence describing over 19,000 gene models for the axolotl. Gene models stop assembling either due to localised low coverage in the genomic reads, or the presence of repeats. We validate our observations by comparison with previously published axolotl bacterial artificial chromosome (BAC) sequences. In addition we analysed axolotl intron length, intron-exon structure, repeat content and synteny. These gene-models, sequences and annotations are freely available for download from https://tinyurl.com/y8gydc6n. The software pipeline including a docker image is available from https://github.com/LooseLab/iterassemble. These methods will increase the value of low coverage sequencing of understudied model systems.

## Introduction

The genomes of salamanders (urodele amphibians) are amongst the largest known (Straus 1971; Keinath et al. 2015; Voss et al. 2011). To date, no salamander genome has been sequenced to completion and only limited genomic data are available (Keinath et al. 2015; Habermann et al. 2004; Smith et al. 2009). Yet this group of organisms are of great interest, not only due to their large genomes, but also their evolutionary history, mechanisms of development and ability to regenerate (Haas and Whited 2017; Swiers et al. 2010; Chatfield et al. 2014). EST sequences and a number of transcriptomes have been generated for *Ambystoma mexicanum* (hereafter the axolotl), covering a variety of developmental timepoints and different stages of regeneration (Habermann et al. 2004; Putta et al. 2004; Evans et al. 2014; Stewart et al. 2013; Jiang et al. 2017; Bryant et al. 2017). Even so, studies are limited by the lack of genome sequence. This makes phylogenetic comparisons difficult and renders many experimental approaches intractable in axolotls as a system. For example, the use of morpholinos, transgenesis and CRISPR-Cas9 for the study of axolotl development and regeneration have all required targeted cloning approaches and Sanger sequencing, or have focussed on transcript sequences alone (Chatfield et al. 2014; Swiers et al. 2010; Sobkow et al. 2006; Flowers et al. 2014; Fei et al. 2014).

The rapid development of sequencing technology has enabled an increased reliance on genomic sequence for studies in model systems. The emergence of long reads, optical mapping and chromatin linkage studies can provide genomic data from more species than ever before (Bickhart et al. 2017). Yet these approaches can be costly and the difficulty of sequencing and assembly scale non-linearly with genome size (Bradnam et al. 2013; Simpson and Pop 2015). Genomes are complex and difficult to assemble for many reasons. These can include GC bias, highly repetitive sequences, sheer scale or even financial limitation. Recently, Keinath and colleagues generated 20x short read Illumina coverage of the axolotl genome (Keinath et al. 2015). In this case, the size of the axolotl genome, combined with the limited coverage, precluded the generation of a useful assembly.

Thus, to date, a significantly detailed genomic sequence has not been generated for a urodele amphibian. Limited sequencing of bacterial artificial chromosomes (BACs) have provided 24 sequences covering less than 0.01% of the axolotl genome, which is estimated to be 30Gb, about 10 times the size of human (Smith et al. 2009). One of the key challenges for a whole genome assembly of this organism is dealing with repeat sequences. Surveys of plethodontid salamanders suggest the most abundant to be members of the LTR/Gypsy retrotransposon superfamily, with repeat lengths of the order of 7 kb, and so likely intractable for assembly using short read approaches (Sun et al. 2012; Sun and Mueller 2014). In total, repeat sequences are estimated to make up as much as 60% of the axolotl genome (Keinath et al. 2015; Smith et al. 2009).

The axolotl genome is an intriguing candidate for sequencing with long read approaches, especially those able to generate read lengths significantly exceeding the length of typical repeats (Jain et al. 2017). However, given the availability of existing whole genome sequence data (Keinath et al. 2015), we reasoned that targeted local assembly around known protein-coding sequences could generate useful data characterising genic regions from the axolotl. These data could identify gene models, intron-exon structures, promoter sequences and more. These resources would be useful for those seeking to exploit common techniques to manipulate gene expression including targeted morpholinos, CRISPR-Cas9 genome editing and even ChIP-Seq.

Borrowing from the ideas of genome walking used in the laboratory (Leoni et al. 2011), we have developed a Virtual Genome Walking (VGW) pipeline which uses localised mapping to known transcripts to extend into surrounding genomic regions. This pipeline is exon-intron aware and allows the input transcripts to be split into exons. Our methodology is similar to previous technologies designed to fill gaps within genome assemblies (such as Gapfiller and IMAGE) (Boetzer and Pirovano 2012; Tsai et al. 2010), assemble flanking data (GenSeed, Tracembler) (Alves et al. 2016; Dong et al. 2007) and assemble exons (Lamichhaney et al. 2012). A similar protocol has been independently proposed by Aluome and colleagues, although no pipeline was made available (Aluome et al. 2016). Our approach is optimised to handle extremely large genome read sets, assemble both flanking and intronic sequence, and continue walking into unassembled genome space iteratively. We have applied this approach to the 20x coverage illumina data from Keinath et al, generating almost 1 Gb of assembled sequence data from the axolotl genome. We validated these assemblies against previously sequenced BACs and find our genome walked fragments closely match with 98.8% identity. Further, we compare the resulting gene models with human and find conservation of exon/exon boundaries within coding sequences as expected.

In total we have generated 19,802 gene models for the axolotl providing the largest assembled set of sequences to date. These sequences, equivalent to a sequenced BAC library, enable the inspection of more of the axolotl genome than previously possible. We note that our completely assembled fraction still represents only 3% of the total axolotl genome, but is equivalent to over 30% of a human genome and is a 300 fold increase in the amount of assembled genomic data available to date. The complete dataset comprising fasta scaffolds, input transcripts and annotations are available for download from figshare (https://tinyurl.com/y8gydc6n). The VGW pipeline is available to download from GitHub (https://github.com/LooseLab/iterassemble). Importantly, the methods we describe can be applied to any model organism with an existing transcriptome and low coverage genome reads.

## Results

### Input Transcriptome

Our previously derived transcriptome dataset (Evans et al. 2014) was re-assembled using CLC, cd-hit and cap3 to generate 646,790 transcripts (see methods, Figure S1, Table 1). Using this entire dataset as input to our VGW pipeline would result in assembling many duplicate genomic loci due to the presence of splice variants or partial transcripts. To avoid this, we identified a single transcript representing each protein-coding gene within our dataset. Transdecoder was used to detect open reading frames (ORFs) which were clustered and annotated into 23,047 protein-coding cDNAs (see methods, Table 1, Figure S1).

**Table 1.** Assembly metrics for our whole transcriptome and final annotated collection of cDNA sequences.

|  | Number | Min Length | Max Length | Mean | N50 | Total (Mbp) |
| --- | --- | --- | --- | --- | --- | --- |
| Assembled cDNA | 646,790 | 176 | 51,730 | 621 | 881 | 402 |
| Annotated cDNA | 23,047 | 297 | 51,730 | 2,325 | 4,121 | 53 |

### Genomic Read Preparation

Genomic reads for mapping to the transcriptome were publically available (Keinath et al. 2015). These data contain over 3 billion 2 x 100 base reads equivalent to 20x coverage of the axolotl genome. Keinath et al were unable to assemble these data using conventional methods, but could estimate the total repeat content of the axolotl genome at between 12-20 gigabases (Keinath et al. 2015). These repeated sequences are likely to hinder both genome wide assembly, and local assemblies around transcripts, particularly when using short reads (Simpson and Pop 2015). Mapping reads to a representative BAC (JF490016) reveals coverage depth exceeding 1,000,000x over some repeats (Figure 1A*i*). Using Khmer (Crusoe et al. 2015), we calculated the median coverage of each read based on 31-mers, discarding paired reads with median coverage less than 1 or greater than 40 (47.8% of the read pairs). A further 8,957,820 (0.3%) read pairs were removed as one sequence contained an ambiguous nucleotide (‘N’). The resultant mapping of this repeat depleted read set to the same BAC demonstrates fold reduction in coverage of up to 70x (Fig 1A*ii*). Even with this reduction in complexity, read mapping was a significant bottleneck to our pipeline (see methods). To resolve this, we indexed the reads rather than the reference and used bwa fastmap to extract super-maximal exact matches (SMEM) (Li 2013). This provides a speed improvement as expected; mapping the repeat-depleted reads to all 24 BACs takes 5 hours with BWA MEM but only 25 minutes with BWA fastmap (30 cores, Intel(R) Xeon(R) CPU E5-2683 v4 @ 2.10GHz). A further advantage of this approach is the increased stringency of mapping, resulting in further reductions (up to 20,000 fold) in repeat coverage with respect to the reference BACs (Fig 1A *iii*).

**Figure 1.**
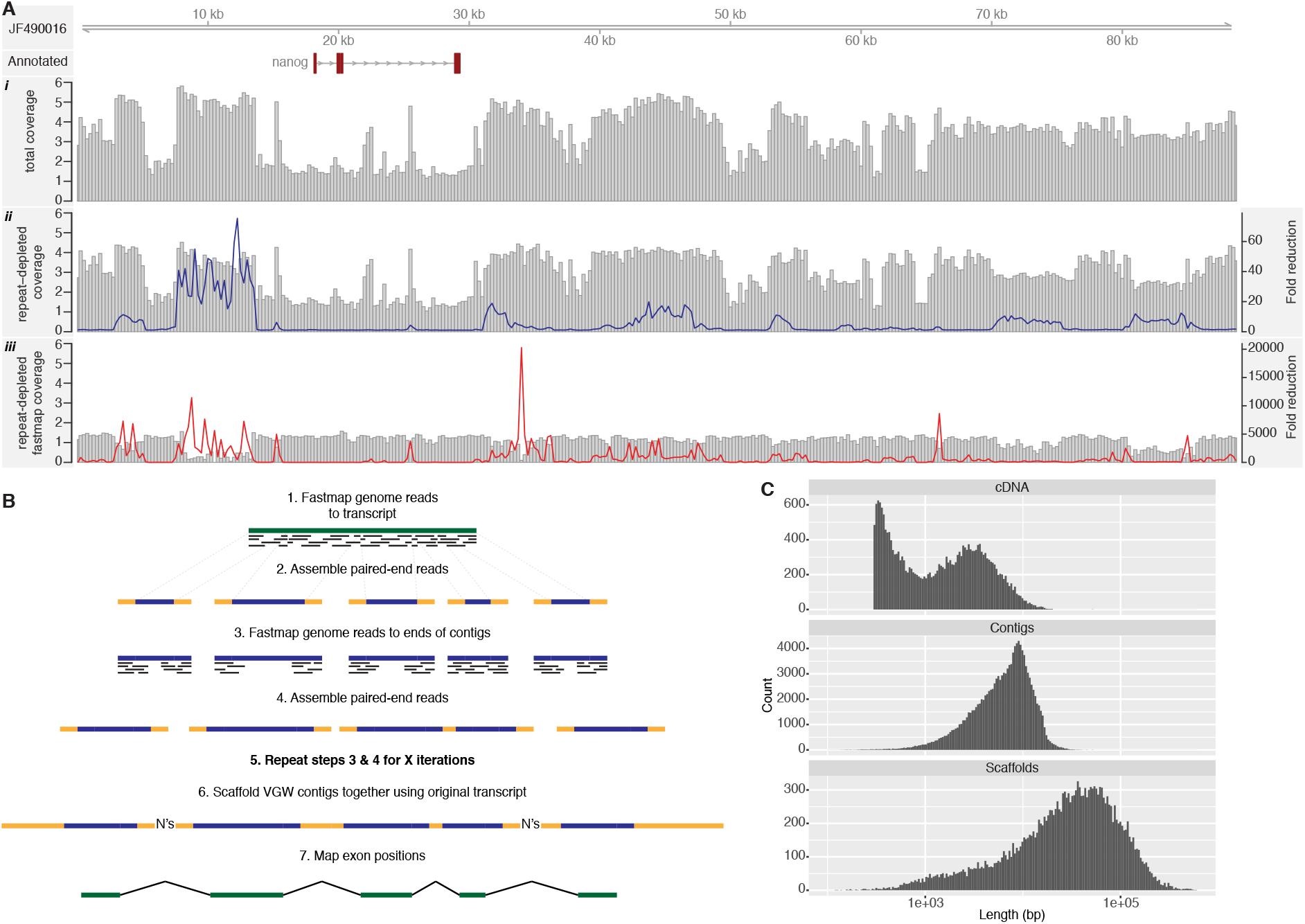
A) BAC JF490016 is shown with the read coverage from all reads (i), the repeat depleted reads (ii), and those that map using fastmap (iii) on a log_10_ scale. The line graphs show the fold reduction in coverage after repeat depletion (blue) and after mapping with fastmap (red). B) Outline of the VGW process. C) Length histogram of the input cDNAs, VGW contigs and scaffolds on a log scale.

### Virtual Genome Walking

We used these repeat depleted genome reads as the input to the VGW pipeline against the unique transcript collection and ran 30 iterations (Figure 1B, Figure S2), generating 22,794 initial genomic scaffolds. A small subset of input transcripts failed to generate any genomic assembly, due to insufficient repeat-depleted reads mapping with a 40 base SMEM. For some of these, no reads mapped to the transcript and we cannot exclude the possibility that these are non-axolotl contaminants. Each genomic scaffold was processed by mapping the input cDNA using GMAP (Wu and Watanabe 2005) and removing any unmapped contigs. Using the newly available intron sequences we were able to identify redundant transcripts and merge the resulting VGW scaffolds (see methods). After processing, our final VGW dataset comprised 19,802 scaffolds in 128,833 contigs (Table 2, Figure 1C). The longest scaffold, at 585,289bp, is derived from a 13,848bp transcript and consists of 79 exons mapped over 63 contigs and is likely orthologous to the human VPS13D gene. Our contig N50 at 9,306bp compares favourably with Illumina only whole genome assemblies from other large genomes (Zimin et al. 2014), although our contigs only include genic regions. The scaffold N50, at 82,884bp, is effectively equivalent to a partially sequenced BAC library.

**Table 2.** The VGW results. Scaffold metrics exclude the artificially inserted 500 ‘N’ gaps between contigs.

|  | Number | Min Length | Max Length | Mean | N50 | Total (Mbp) |
| --- | --- | --- | --- | --- | --- | --- |
| Contigs | 128,833 | 100 | 52,850 | 7,313 | 9,306 | 942 |
| Scaffolds | 19,802 | 119 | 585,289 | 47,578 | 82,884 | 942 |

We arbitrarily stopped VGW after 30 iterations to investigate how the algorithm had performed. Given the prevalence of high copy long repeats in the axolotl genome, we hypothesised that repeats would be the most common cause of VGW failing to extend a contig. Alternatively, since the genomic reads represent a depth of only 20X, insufficient coverage of specific regions might prevent a contig from extending. To investigate this we ran an additional iteration to identify those transcripts that continued to extend (see methods). This revealed that 14,179/22,794 (62.2%) transcripts that began extending were still being processed by iteration 30. However, the majority of contigs within each extending transcript had stopped with only 30,420 (23.6%) contigs still extending by iteration 30. To distinguish between contigs stopped by low coverage compared with repeats, we calculated the mode depth of coverage at the ends of extending contigs compared with the depth of coverage at those which had stopped (Figure 2A - see methods). As anticipated, in the majority of cases VGW stopped extending due to repeat sequences (76.3%, Figure 2B).

**Figure 2.**
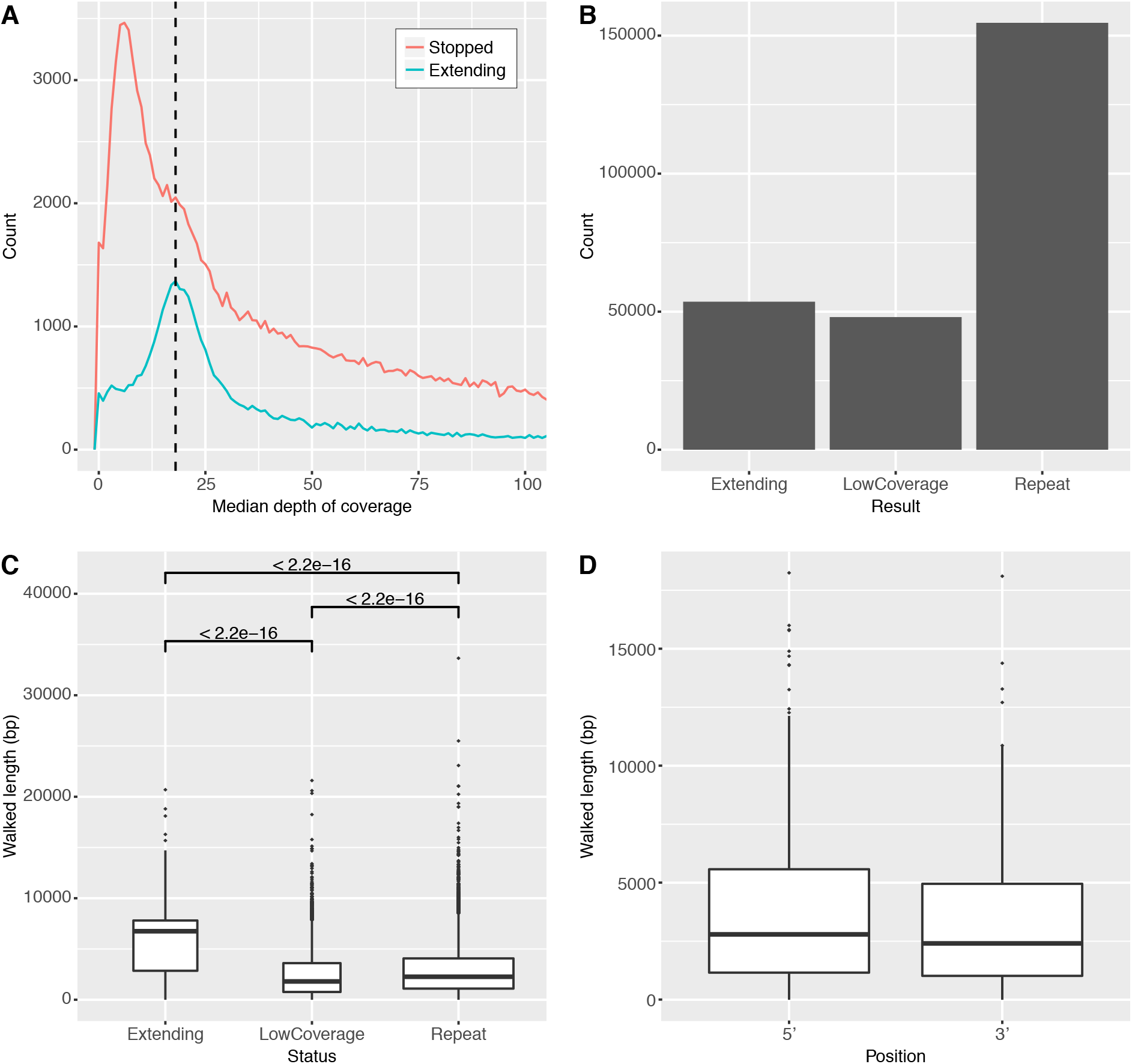
Most contigs stopped extending due to repeats. A) shows the median depth of coverage over the last 60bp of each contig, the results are separated based on whether the end of that contig continues to extend in iteration 31. The mode depth for extending contigs (dashed line) is used to divide the others into low coverage and repeats. B) Most contigs stopped extending because of repeats, defined by a high median coverage. C) The length walked from the nearest mapped exon is shown according to the contig status. D) The length walked from the outermost exons is shown, irrespective of whether the contig had stopped extending.

By mapping transcripts back to the genome fragments, we were able to determine the length walked from the closest exon (Figure 2C). Contigs still extending had walked significantly further from the closest exon than those stopped due to repeats or low coverage. However, a subset of contigs marked as extending are shorter than expected. Presumably these contigs either fluctuate in length from iteration to iteration or are only extending by a small number of bases, both of which we have observed on inspection. The median length walked from the nearest exon for all contigs still extending is 6,717bp, thus VGW is extending the end of each contig by approximately 220bp each iteration. We also examined the distances walked from the outermost exons of each gene model, suggesting that we were able to assemble 2,607bp on average (median) of flanking DNA either side of each gene (Figure 2D). Given this, we assume that the majority of promoter sequences will be included within the VGW scaffolds as human promoter binding sites are within 30-300bp of the transcription start site (Smale and Kadonaga 2003; Koudritsky and Domany 2008).

### Classifying Repeats

Given the prevalence of repeat sequences throughout our data, we generated a repeat library based on all available axolotl genomic data, including the available BAC sequences (see methods). Using this library of 2,103 classified repeats, RepeatMasker masked 34.90% of the VGW scaffolds. The majority of masked repeats were classified as unknown (19.96%), but 7.98% were identified as LINE and 5.27% as LTR elements. The mean length of masked repeat sequence was 225 bases, shorter than the mean length of masked repeats on the BACs (283 bases), but longer than the peaks of high coverage (Figure 3A). The longest repeat masked in the VGW scaffolds is 5,695bp (unknown class), suggesting that in some cases we have genome walked further across, presumably diverged, repeats than anticipated (Figure 3B). Furthermore, only 12.4% of masked repeats were located within 60bp of the end of a contig, suggesting that the majority of repeats masked were walked through to completion. This may be due to the high stringency of BWA fastmap negating problems of high coverage in divergent repeats, as suggested in Figure 1A.

**Figure 3.**
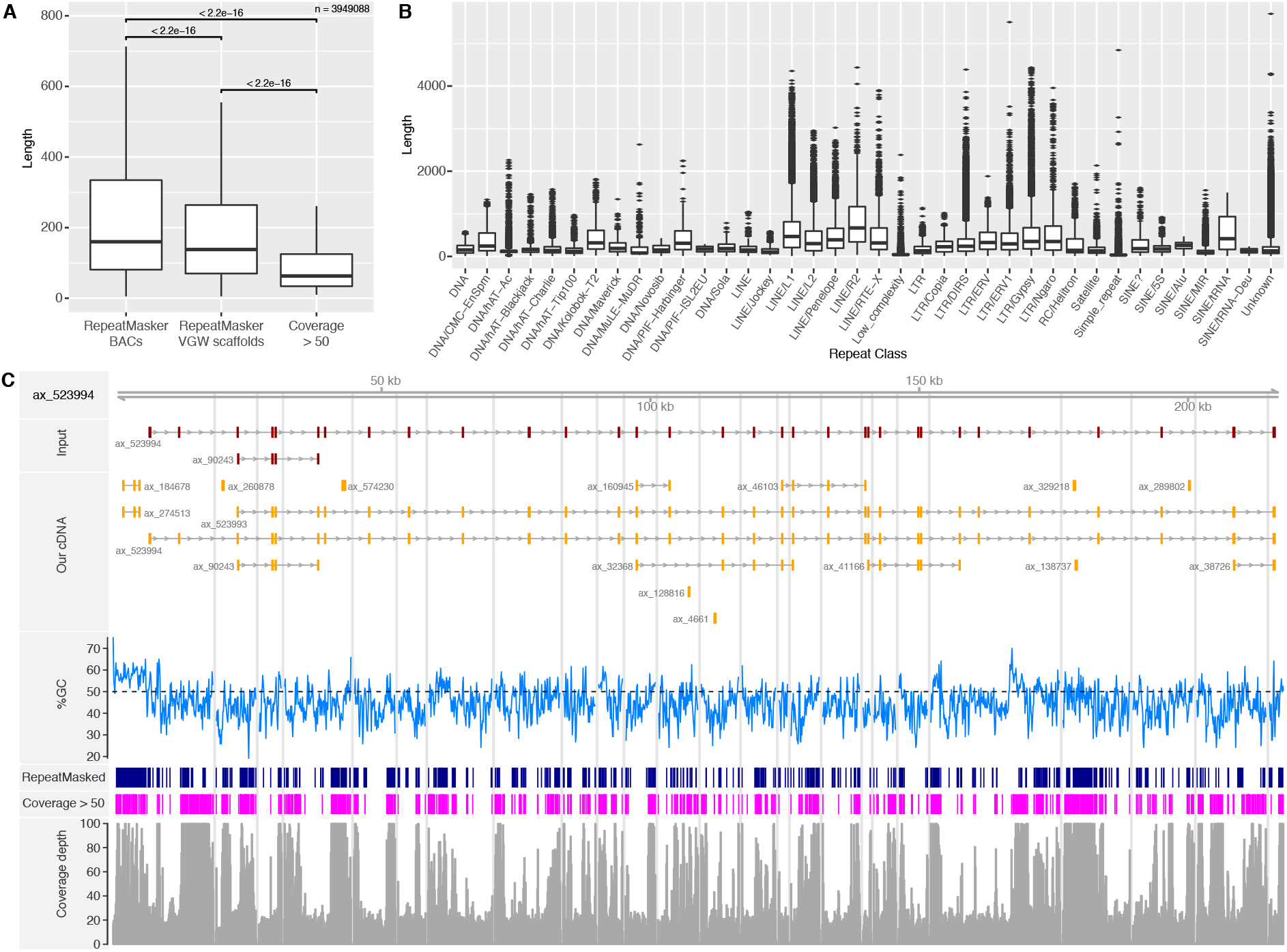
Repeats. A) Boxplot showing the difference in repeat lengths between those identified by RepeatMasker on either the BACs or VGW scaffolds, and those with coverage greater than 50X. B) The length of repeats identified within the VGW scaffolds, separated by repeat class. Only repeat classes that were identified more than 500 times are shown. C) The VGW output for ax_523994, this gene has the longest repeat identified of 5,695bp. The repeats masked are shown in dark blue, the regions with coverage greater than 50 are shown in magenta. The coverage depth, up to a maximum of 100, is shown at the bottom. Contig breaks are shown by the grey bars.

The most common repeat identified within the VGW scaffolds was an unknown repeat named ‘rnd-1_family-16’, which was masked 15,251 times with a consensus sequence of only 155 bases. We believe that this repeat is likely derived from a small non-coding RNA or retroposon similar to other repeats that have previously been observed in salamanders (Batistoni et al. 1995). The next most common repeat outside of the unknown class was a LINE/L2 element (1,488 bp consensus), not an LTR/Gypsy repeat as suggested by Sun et al (Sun et al. 2012), although this does agree with the observations by Keinath et al. on repeat distribution (Keinath et al. 2015). Figure 3C shows ax_523994, which contains the longest masked repeat; regions with coverage greater than 50 are also shown. Peaks of coverage, and masked repeats, often appear at the ends of contigs, suggesting these repeats caused the VGW to stop extending.

### Assessing scaffold quality

Before analysing the VGW scaffolds in detail, we needed to be confident that our assembly is of high quality. To do this, we compared the VGW scaffolds against the 24 publicly available BAC sequences (Smith et al. 2009). We were able to identify 16 BACs with corresponding VGW scaffolds (see methods, table S1). The VGW scaffolds mapped with a mean identity of 98.8%, the same mean identity as the cDNA sequences map to the BACs, suggesting the VGW scaffolds are similar in quality to transcripts assembled using conventional methods. Fourteen VGW scaffolds mapped contiguously to the BACs, the longest of which is illustrated in Figure 4A. This VGW scaffold is comprised of only two contigs and contains a complete gene model annotated as collagen type 1 alpha 2 chain (COL1A2). There are 52 mapped exons in this gene; the longest intron is 5,064 bp and has been completely bridged by VGW. Furthermore, we have managed to walk across a previously un-assembled gap in the reference BAC. This example also demonstrates that the VGW contigs stopped extending at both regions of low coverage and high-copy repeats. Interestingly, the 5’ contig could not be extended beyond the same point as a gap in the BAC suggesting that irrespective of method, short reads are incapable of assembling this genomic region.

**Figure 4.**
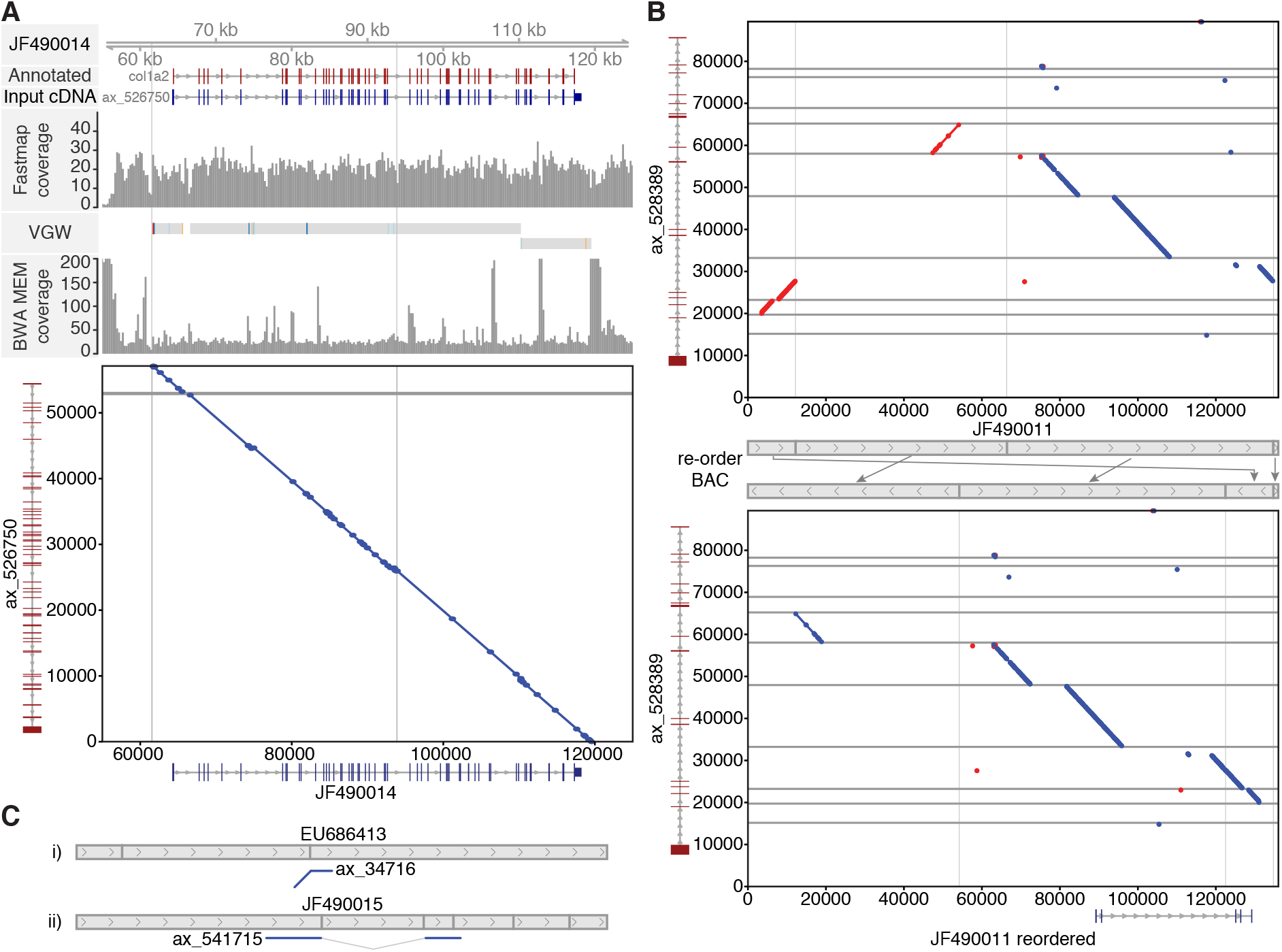
Comparing the VGW scaffolds and BAC sequences. A) A high quality, long VGW scaffold maps to BAC JF490014. Horizontal and vertical grey bars represent gaps within the VGW and BAC assembly respectively. Exact matches of 20bp or more are shown in red and blue in the dot-plot, representing the forward and reverse direction respectively. B) The dot-plot shows that some of the JF490011 BAC fragments are in a different orientation to the VGW scaffold. After re-orientating the BAC fragments according to our assembled scaffold, we are not only able to map the transcript but also show how we walked across one of the BAC gaps. C) shows cartoon examples of two scaffolds that are contiguous with the BAC, yet the BACs are most likely mis-assembled. Dot-plots for these two comparisons are shown in Figure S3.

There were two non-contiguous scaffolds that mapped to fragmented BACs, one of which is illustrated in Figure 4B. Here the VGW scaffold maps contiguously to each fragment but the fragments are incorrectly ordered and orientated with respect to the VGW scaffold. One contig maps to two non-contiguous BAC fragments, suggesting that the BAC may be incorrectly assembled. Although the transcript could be aligned to the BAC with 99.4% identity, GMAP was unable to map the gene model with correct intron-exon boundaries. After re-orienting and ordering the BAC fragments based on the VGW scaffold, GMAP was able to map some of the exons. Originally this BAC was not annotated with a gene model, presumably due to the mis-assembly. The other VGW scaffold that appeared noncontiguous also suggested that it was due an incorrectly assembled BAC (Table S1). Furthermore, two other VGW scaffolds, although contiguous, suggested that their respective BACs were incorrectly assembled (Figure 4C, Figure S3). One of these shows the VGW contig extending beyond the BAC fragment, yet that fragment is positioned within the middle of the BAC. In the other case, a single VGW contig skips a central fragment suggesting the BAC has been incorrectly ordered.

Furthermore, it should be noted that not all contigs within these scaffolds are able to map to the BACs. These additional contigs are derived from exons that are beyond the BAC sequence. In ax_528389 (Figure 4B) there are 7 additional exons upstream of the reordered BAC, and 2 exons downstream, all of which have generated genomic contigs. Indeed, across the 16 VGW scaffolds being compared to the BACs, only 7 contain the same number of exons as on the BAC (Table S1). This highlights how VGW is able to assemble and scaffold genomic fragments containing a single gene, irrespective of the distance between exons.

### Examining Introns

To analyse intron lengths within our new gene models, we analysed transcripts that mapped in a single path to their corresponding VGW scaffold. Of these, 15,364/18,448 paths had more than one exon mapped to the genome. We removed a further 184 gene models from our analysis as an intron contained multiple contigs, and so the length was unreliable. The 15,180 gene models remaining contained 134,002 introns, of which 35,300 (26.34%) were completely assembled by VGW. A total of 689 gene models (4.54%) were mapped to a single contig, with 378 of these containing only a single intron. As expected, the introns we are able to completely assemble are significantly shorter than those that remain unassembled (Figure 5A, Figure S4). This suggests that high-copy, conserved repeats that we are unable to walk across are present within long intron sequences in axolotl. Figure S5 shows two examples of VGW fragments, the first shows one of the longest introns we are able to assemble (16,818bp, ax_530571 orthologous to human LRIG3). The other shows the longest completely assembled gene model that maps to a single contig of 43,023bp (ax_572447, orthologous to human MYT1).

**Figure 5.**
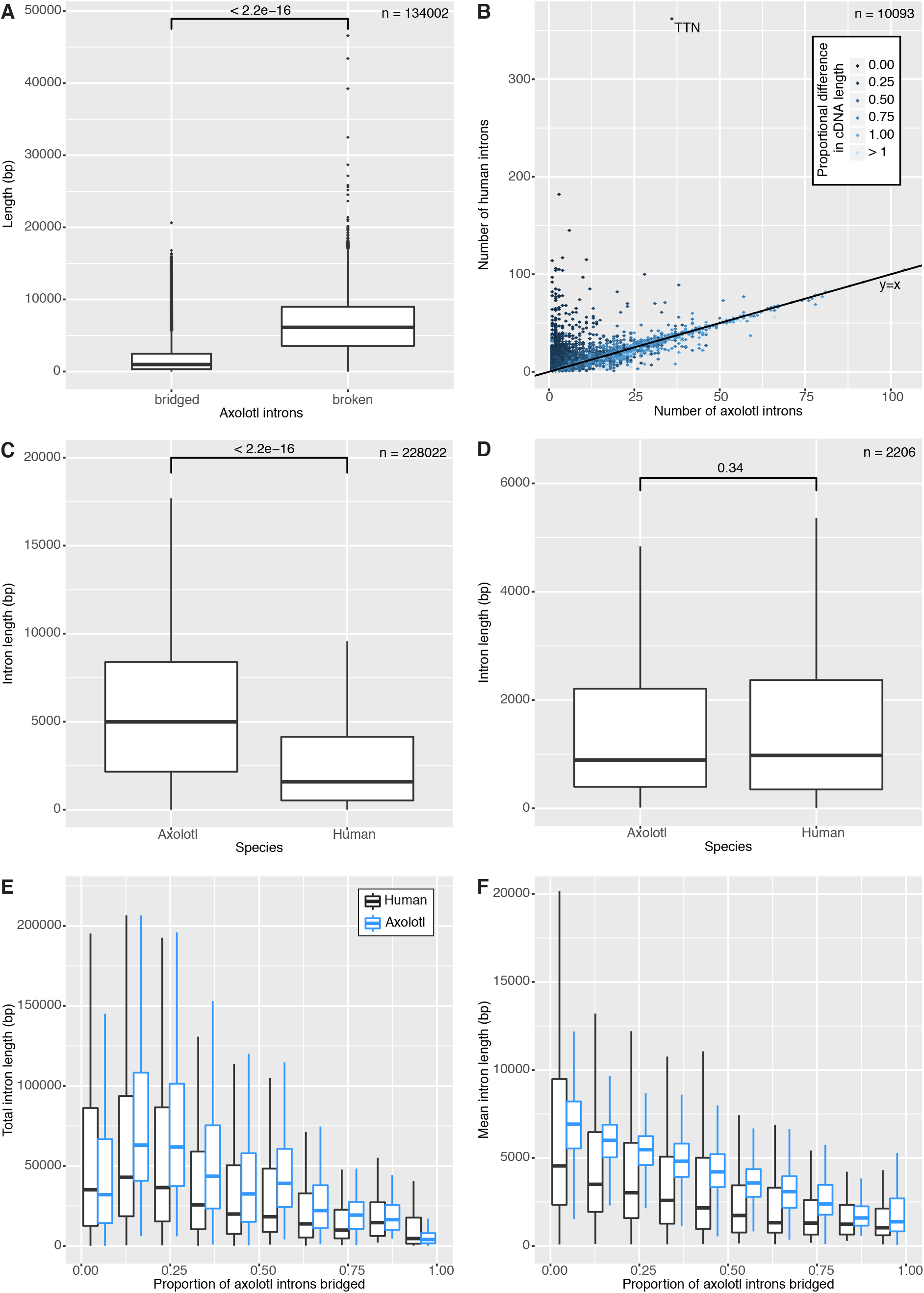
The Axolotl introns. A) boxplot of axolotl intron lengths comparing those we can bridge, and those we cannot. B) Number of introns in axolotl and human, plot is coloured by the ratio of axolotl to human CDS lengths. C) Intron length distributions in axolotl and human. D) Intron length distribution in axolotl and human for completely bridged genes. E) Total intron length per proportion bridged in axolotl (blue) and human (black). F) Mean intron length per proportion bridged.

Based on analysis of relatively few BAC sequences, Smith et al proposed that axolotl introns are approximately 10 times larger than in other vertebrates, consistent with the overall genome size expansion (Smith et al. 2009). To investigate this, we aligned the 15,180 transcripts described previously against human protein-coding genes using BLASTn. 12,604 genes matched with an e-value of less than 1e-10, of which 10,093 were unique ensembl IDs (see methods). This revealed that 3,857 (38.21%) axolotl transcripts have the same number of introns as in human, and that 8,613 (85.34%) show a small difference of +/-5 introns (Figure 5B). As expected from a potentially incomplete transcriptome assembly, there are many genes with more introns in human than axolotl, for example titin. Furthermore, the exon boundaries appeared conserved with human, with over 65% of the axolotl boundaries present within 10bp of the human exon-intron boundary (see methods, Figure S6). Although this appears less conserved than *Xenopus*, we believe the difference is due to small inaccuracies in GMAP defining the exon positions and not a difference in the underlying biology.

Although many axolotl introns are incomplete, they are still larger than the equivalent human introns on average (median 3x larger) (Figure 5C). However, fully bridged genes show no significant difference in intron length (Figure 5D). Indeed shorter genes that we are able to bridge in VGW also tend to be shorter in the human genome (Figure 5E and F). There are some cases where gene models are significantly expanded with respect to the human genome. For example, ax_517324 has a scaffold length in excess of 450kb even though almost none of the introns are bridged (Figure S7). Yet its ortholog in human (ENSG00000123384, LRP1) only covers 100kb. However, ax_576019, for which every intron is completely bridged, has a total intron size of only 3kb. The ortholog in humans (ENSG00000143333, RGS16) differs in total intron size by only 49bp, although the individual introns do vary. Both of these example genes contain the same number of exons in Axolotl as in Human. Overall, our results suggest that intron size is gene dependent and that the genome size expansion is not uniform across the genome.

### Transcript mappings

By mapping additional RNA-seq datasets to each VGW scaffold, we are able to distinguish transcript variants from gene copies (Bryant et al. 2017; Jiang et al. 2017). An example gene with transcript variants including both alternative exon usage and differing splice sites is shown in Figure S8. Similarly, genes thought to have multiple copies in axolotl can now be confirmed using genomic sequence, such as Nodal and Brachyury (Figure S9 and Figure S10)(Swiers et al. 2010). As only one copy of each gene is present in humans, both axolotl Nodal genes are annotated as the same gene in the Jiang dataset, as are both copies of Brachyury. This demonstrates the increase in resolution that can be obtained by mapping against VGW scaffolds.

A subset of transcripts are able to map to more than one VGW scaffold. In the majority of cases, these mappings were low identity and likely derived from simple repeats or regions of low complexity. However in several cases one or more transcripts mapped with high identity outside of the known exon positions. Furthermore, in 248 instances the two VGW scaffolds shared identity across this region, suggesting these genomic fragments might be linked. The most likely cause is fragmented transcripts, as poorly expressed genes may not have been completely assembled in the transcriptome. To assess this, we analysed the Jiang and Bryant datasets (Jiang et al. 2017; Bryant et al. 2017), looking for transcripts which overlapped exons on both VGW scaffolds. This identified 210 pairs that were likely derived from the same gene, demonstrating that we have walked across introns that we did not know existed. For example, Figure S11 shows a single scaffold derived by merging three VGW scaffolds to which 6 of our input cDNAs map. In the Jiang dataset, there is a single transcript that maps to all of the identified exons, suggesting that this is one gene.

There were still 38 pairs of scaffolds that could not be joined by a single Jiang or Bryant transcript. To analyse these further we identified their most likely human ortholog using BLASTn. Only 20 pairs found a human ortholog for both axolotl transcripts with an e-value less than 1e-10. Exactly half of these were also derived from fragmented transcripts from a single gene, identified either through non-overlapping BLASTn HSPs to the same human gene, or manual inspection. Of the remaining 10 pairs of linked VGW scaffolds, one appears to be a repeat as both transcripts map within repeat masked sequence. Two of the pairs may be mis-assemblies, as the linked contigs contain only a single exon that is not represented in either the Bryant or Jiang transcripts. Furthermore both of these axolotl genes contain an additional exon compared to their human ortholog. This could therefore be a case of transcript mis-assembly leading to an incorrect VGW scaffold that appears to contain two genes. This leaves 7 pairs of axolotl scaffolds that appear to be syntenic, as they contain two distinct genes that we have walked between. This is corroborated by the human orthologs, 5 of which share synteny, indeed they are the closest genes to one another in the human genome (Table 3). The mean intergene distance between these human genes is 1,130bp, although 3 genes overlap and therefore have a distance of 0. The mean intergene distance in axolotl is 1,722bp, and is not significantly different to the human distance (p = 0.0625; Wilcoxon signed-rank test). The remaining two genes are not syntenic in Human, Chicken or Xenopus; without further validation of these VGW scaffolds it is not possible to know if these are genuinely syntenic in axolotl.

**Table 3.**
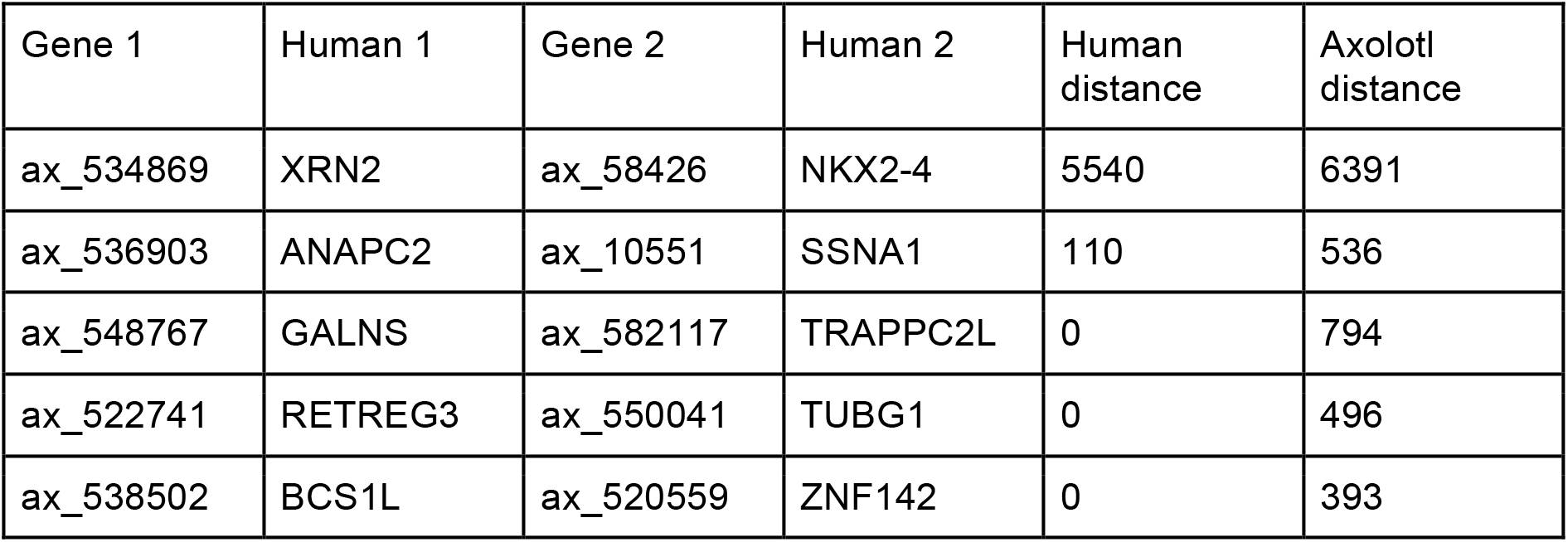
Syntenic genes that we have walked between. Distances of 0 are shown for genes with overlapping exons.

One example of a syntenic gene is shown in Figure S12, the VGW scaffolds of ax_538502 and ax_520559 are assembled together using CAP3. There are two distinct genes mapped to this single scaffold, orthologous to BCS1L and ZNF142 respectively. Both axolotl genes share the same number of exons as in human, and are only separated by 393bp. This further suggests that the axolotl genome size increase is not uniform.

### Chromosomes 13 and 14

Finally, we looked at whether the chromosome capture sequencing of chromosomes 13 and 14 could be used to distinguish which genes are situated on these chromosomes (Keinath et al. 2015). To do this we first associated 615 VGW fragments with the known linkage groups (LG) based on at least two primers aligning to our transcripts (Voss et al. 2011). We then compared the coverage depth across exons of the 40 linked genes on AM13 (LG15 and LG17) and the 19 AM14 (LG14) genes with those on the other linkage groups. Unfortunately, the appropriate SRA files were not annotated with chromosome of origin. Therefore to distinguish from which chromosome the reads were derived, we first compared each uploaded SRA file and grouped them accordingly (Figure S13A). We then re-mapped the data using these grouped files to improve coverage. Although the chromosome capture sequencing was at a low coverage, we were able to enrich for genes present in the chromosome 13 and 14 linkage groups (Figure S13B-F).

We then mapped these reads to all VGW scaffolds, and again calculated the coverage depth across the exon positions. By using the parameters identified for the linkage group genes (Figure S13), we were able to isolate 1,708 axolotl VGW scaffolds likely to be on AM13, of which 596 transcripts had a blast result in human (e-value < 1e-10, 546 unique Ensembl IDs). We were able to isolate 1,397 VGW scaffolds likely to be on AM14, corresponding to 399 human genes, of which 368 are unique. We note that the original number of axolotl genes isolated is higher than expected for the two smallest chromosomes considering that if all 14 chromosomes had an equal number of genes we would only see 1,414/19,802 genes per chromosome. Furthermore, there are a large number of genes with no BLASTn alignment to human protein-coding genes, which suggests the outstanding axolotl transcripts may be transposases present on more than one chromosome. Indeed, 656 of the isolated axolotl transcripts are shared between AM13 and AM14, only 21 of which have a human BLASTn match.

By analysing the location of the human orthologs to the unique AM13 and AM14 axolotl genes we are able to compare synteny between axolotl and human. The AM13 genes were associated with human chromosomes 17, 1 and 6 while the axolotl AM14 genes were associated with chromosomes 14 and 15 (Figure 6). The regions on these human chromosomes are highly syntenic with the chicken chromosomes 26, 5 and 27; all of which were previously highlighted as being syntenic with axolotl chromosomes 13 and 14 (Keinath et al. 2015). Therefore, the chromosome capture sequencing reads can be used to classify axolotl VGW scaffolds consistent with direct assembly methods.

**Figure 6.**
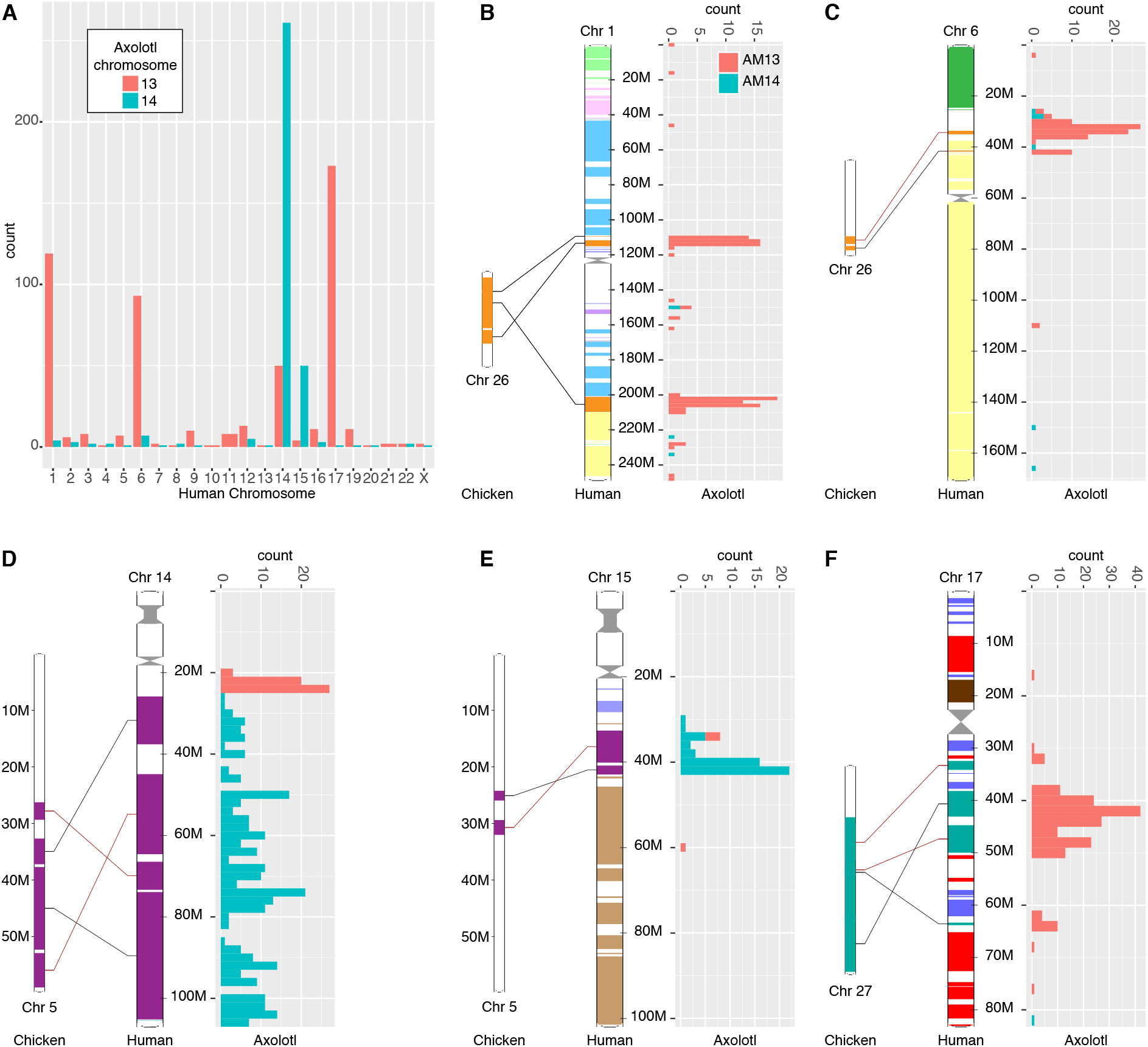
Synteny between axolotl, human and chicken. A) The number of axolotl genes on AM13 and AM14 orthologous to genes on human chromosomes. The location of the orthologous genes are shown, along with synteny between that region in human and chicken (Ensembl) for human chr1 (B), chr6 (C), chr14 (D), chr15 (E) and chr17 (F).

### Comparison to whole genome assembly

While writing up this work, a whole genome assembly utilising Keinath et al’s previous paired-end reads and a new mate-pair library was released as an unpublished queryable BLAST database (www.ambystoma.org). The assembly consists of 21Gb in 21 million scaffolds with an N50 of 27,236bp. Although a complete comparison is not possible due to limited access, we used BLAST to investigate 200 randomly chosen VGW scaffolds and extracted all sequences with an e-value of 0 and bitscore greater than 1000. We specified this level of stringency to limit spurious matches caused by repeats. Nevertheless, 185 of our VGW scaffolds extracted 3,531 whole genome scaffolds. We identified and removed 2,040 of these potential BLAST matches as mapping to repeats because they either mapped to multiple VGW scaffolds or contained overlapping HSPs to the same VGW scaffold. To better understand the remaining 1,491 whole genome scaffolds (Table 4) we split the sequences on gaps of at least 100 Ns and identified these as contigs. Considered either as scaffolds or contigs, the mean and N50 values are shorter than our VGW sequences. Over these genic regions the transcriptome provides more information than the mate-pair library alone. 24% of our VGW contigs are represented by more than one whole genome scaffold, as is apparent from the large number retrieved by BLAST.

An example case is shown in Figure S14, where a VGW scaffold of 117kb is aligned against a whole genome scaffold of 74kb. The putative exon positions were identified by mapping the VGW transcript against both scaffolds. The whole genome scaffold is contiguous with our assembly, providing an independent validation of our results. Indeed, across the 185 VGW scaffolds with a BLAST hit, 12.6% of the 1,187 contig breaks were supported by whole genome scaffolds. However, only 19 (1.6%) were completely assembled without gaps of more than 100 Ns. Figure S14 also shows that most of the gaps in the VGW scaffold are also represented with Ns in the whole genome scaffold. This suggests that the axolotl repeats that stop VGW cannot be resolved using mate-pair assembly alone.

**Table 4.** The assembly metrics for 200 random VGW scaffolds and their corresponding whole genome (WG) scaffolds.

|  | Number | Min | Max | Mean | N50 |
| --- | --- | --- | --- | --- | --- |
| VGW - scaffolds | 200 | 638 | 320,175 | 50,759 | 86,249 |
| VGW - contigs | 1395 | 144 | 28,192 | 7,277 | 9,146 |
| WG - scaffolds | 1,491 | 565 | 370,994 | 25,104 | 39,788 |
| WG - contigs | 13,671 | 14 | 83,208 | 2,738 | 4,723 |

## Discussion

Here we have demonstrated how low coverage Illumina data can be used to generate gene models even when the raw reads themselves cannot be assembled by conventional methods. Using the 20x Illumina reads generated by Keinath et al. we have been able to assemble almost 1 Gb of the axolotl genome (Keinath et al. 2015). These sequences are not random portions of the genome, rather they represent gene models for over 19,000 genes. This is a unique and invaluable resource for the field that will permit studies on evolutionary genomics and the use of genomic molecular biology methods on a salamander model species. As well as building gene models and assembling intronic sequence, we have been able to walk on average 2kb upstream and downstream of each transcript. In principle this data set should include the majority of promoter sequences for each transcript and enables the use of ChIP-seq methods against promoters in the future.

Using the dataset presented herein we have generated a new axolotl repeat library containing 2,103 consensus sequences. This demonstrates that our VGW algorithm allowed us to walk through an unexpectedly high number of repeats. Although the majority of these were short, we were able to assemble repeats >1kb. In total, over 30% of our VGW scaffolds were identified as being repeat sequence; we anticipate a higher proportion of repeats outside of the genic regions as suggested in Figure 1A.

We have demonstrated that the genome gigantism observed in salamanders does not appear to be uniform across genic regions in the axolotl genome. Indeed, gene model lengths in axolotl correlate with their orthologous lengths in human. Genes that we have completely assembled show no significant increase in overall intron size compared to their human orthologs. Where introns cannot be bridged, the minimal expansion is 3x the equivalent distance in the human genome. The upper estimate on expansion is at least 10x based on the observations of (Voss et al. 2011). Furthermore, in a handful of cases we had successfully walked between axolotl genes, suggesting that intergenic regions are not uniformly expanded either.

We have utilised additional resources built for the axolotl community to map some of our assembled scaffolds to chromosomes 13 and 14. Our approach demonstrates that even low level coverage of individual chromosomes is sufficient for these classifications. Therefore, continued short-read sequencing of chromosome capture, combined with the known linkage map, could allow further classification of both our VGW scaffolds and whole genome assembly scaffolds.

Whilst writing up this work, a new Illumina based assembly employing mate-pair libraries was released. This data set, representing almost 21Gb of the axolotl genome is currently only accessible by blast searching and has a scaffold N50 of 27,236 bases (see www.ambystoma.org). We have tested a subset of these with super contigs released by Voss and colleagues and show that our contiguity agrees well with whole genome assemblies. Interestingly, we find that some regions cannot be assembled through in either data set, confirming that short reads alone will be insufficient to dramatically improve the assembly contiguity. Rather a long read strategy is likely required incorporating either Pacific BioSciences and additional technologies (Bickhart et al. 2017) or ‘ultra-long’ reads from the Oxford Nanopore platform (Jain et al. 2017). It will be interesting to determine if VGW using more comprehensive read sets including mate-pairs will enable higher quality gene models than simple whole genome assembly.

The VGW method itself has general utility for analysing genomes with low coverage read sets available. Whilst cost is not typically a limiting factor for traditional model organisms and human genomes, it is often a concern to those working on less well studied models. Such models are often found to have larger genomes, which substantially increases the cost of sequencing. Such genomes also tend to be repeat rich, requiring elaborate library strategies to resolve on conventional short read platforms. We have shown here that useful genomic data can be recovered in such circumstances from limited coverage of a reference genome Indeed, for many common genomic approaches a whole genome is not absolutely necessary, rather a comprehensive targeted approach can provide as much benefit as the entire genome.

## Methods

### Transcriptome sequencing and Assembly

The axolotl RNA was isolated from oocytes and a range of developmental stages (Evans et al. 2014). Each library was initially assembled using CLC (QIAGEN Bioinformatics) with the default parameters, we then merged each library by clustering with cd-hit and assembling each cluster using CAP3 (Li and Godzik 2006; Huang and Madan 1999). To identify protein coding regions, Transdecoder was first run to identify all long putative open reading frames, these were then blasted against a vertebrate specific protein nr database (Evans et al. 2014; Grabherr et al. 2011). Transdecoder used both datasets to predict the best ORF per starting transcript. In a few minority cases, TransDecoder preferentially chose the longer ORF over one with protein homology, we therefore wrote a custom script to select the ORF with homology in these cases. Redundancy was then removed from the dataset using cd-hit and our custom program which clusters using BLAST, in both cases the longest ORF per cluster was retained. To annotate these non-redundant sequences, we ran them through BLAST2GO which uses protein blast results to describe and assign GO-terms. This step removed a large number of sequences derived from transposases. Our final dataset consisted of 23,047 annotated cDNAs as described in the results section.

### Repeat Depletion

To repeat-deplete the Keinath et al whole genome reads, each read file was run through khmer to calculate the median coverage based on kmers of 31bp (Crusoe et al. 2015). Read pairs were removed if either read had a coverage less than or equal to 1 or greater than 40. We also removed any read pairs which contained an ambiguous nucleotide (‘N’) using a python script available alongside VGW.

### Virtual Genome Walking

The Virtual Genome Walking (VGW) pipeline first divides the input genome read files into multiple sub-files, each of which will be indexed using BWA. Optionally each of these subfiles can be compressed using gzip at the expense of running speed. Once created, the index can be reused indefinitely.

On the first VGW iteration, transcript sequences are mapped to indexed reads using BWA fastmap with a minimum super maximal exact match (SMEM) of 40bp (Li 2013). This permits reads to map to almost all exons. The mapped reads are then extracted into separate files per starting transcript, and assembled using SOAPdenovo2 (k = 63bp) and fermi-lite (https://github.com/lh3/fermi-lite/) with default parameters (Luo et al. 2012). The output from both processes are assembled using CAP3 (-k 0 -p 75 -o 30 -h 80 -f 200 -g 4) with parameters that permit longer overhanging sequences. The original transcript and CAP3 output are aligned using BLASTn with a culling limit of 2 to extract assembled contigs that contain exon sequence. Finally, the extracted reads are mapped using BWA and any contigs that share sufficient paired-end read mappings with the exon-containing contigs are also retained. This ensures that any close neighbour contigs that do not contain exons are extended in the following iterations.

On all following iterations, 600bp at the ends of each contig are mapped to the reads using a longer SMEM of 60bp. As before, SOAPdenovo2 and fermi-lite are used to locally assemble the extracted reads. An additional elongation step is added whereby the short reads mapping to each contig are assembled using CAP3 with default parameters. These contigs are combined with the SOAPdenovo2 and fermi-lite contigs, and the final contigs from the previous iteration prior to assembly with CAP3 (overhang parameters). The exon-containing and potentially neighbouring contigs are identified as before. After each iteration, the maximum contig length and the sum lengths are compared to the previous iteration. If either of these has increased, then another iteration will be run on that transcript. The number of contigs is also compared and if it reaches a set maximum value (by default 500) or triples within a single iteration then that transcript is no longer processed. This ensures that repeat containing contigs do not disrupt the VGW process by assembling sequence from across the genome.

After the final iteration is complete, we first check that each contig appears correctly assembled. The transcript and contigs are aligned by BLASTn, identifying the contigs that contain multiple HSPs to the same exonic sequence. For each of these potentially problematic contigs, all of the previously extracted reads are mapped using BWA. At each position along the contig the coverage of paired end reads mapping with 100% identity is calculated. Any region with a coverage of 0 that is not at the ends of the contig, or has no paired end reads spanning it, is removed to divide the chimeric contig into sub contigs.

Contigs are reverse complemented according to the transcript, and each contig is compared against each other by BLASTn to search for large regions of similarity. Within each group a consensus sequence is derived by aligning the contigs using MAFFT (Katoh et al. 2002). Therefore alternative assemblies of the same locus, which may differ in the length of a repeat, do not appear as contiguous genome fragments.

The re-orientated, consensus derived contigs are ordered according to BLASTn HSPs with the original transcript. Each contig is aligned with the following contig using BLASTn to look for a potential overlap at the respective ends. If there is an overlap, then the contigs are joined using a consensus sequence of the overlap region (MAFFT). Contigs with no perceivable overlap are joined with an arbitrary string of 500 ‘N’s, generating a single genome scaffold per original transcript.

### VGW scaffold processing

Prior to analysis we parsed the VGW scaffolds according to the original transcript. We used GMAP to find exon-intron aware cDNA matches in each VGW scaffold using the starting transcript (Wu and Watanabe 2005). Some contigs did not contain an exon mapping to them, this appeared to be due to closely related gene copies with a shared exon sequence. VGW was walking out from two copies of near identical exons, only one of which could be successfully mapped to by GMAP. We therefore removed these non-mapping contigs, identified through the 500 ‘N’ breaks, from the scaffolds.

To identify redundant VGW scaffolds we re-mapped the transcripts using GMAP to the parsed dataset. Starting from the longest VGW scaffold, we used the GMAP GFF3 file to identify additional transcripts with overlapping exon positions. We then compared the VGW scaffolds derived from these transcripts using BLASTn, so long as all exon positions were within an HSP of at least 95% identity and at least one was positioned within an HSP of 99% identity the scaffolds were merged. We used CAP3 to assemble the contigs from all VGW scaffolds together. For any new contigs that could not be assembled this way, we first checked if they shared identity with any other contigs and if so, used MAFFT to align and generate a consensus sequence. The final collection of contigs were ordered according to how each transcript mapped, generating a single VGW scaffold per axolotl gene.

### Genome Visualization

Genome visualization was done in the R/Bioconductor package GVIZ (Hahne et al. 2013). Exon positions were identified using GMAP. Coverage was determined using all repeat-depleted reads mapped with BWA MEM and samtools depth unless otherwise stated (Li 2013; Li et al. 2009). To import into R, coverage and %GC means were averaged over 20bp windows, they were then visualised in fixed 100bp windows. For the BACs, no prior compression was required and the coverage depths were visualised in 250bp windows. For clarity only GMAP results with more than one exon are displayed for the larger datasets of all cDNAs and the Jiang and Bryant transcripts unless otherwise stated (Jiang et al. 2017; Bryant et al. 2017).

### Extending VGW contigs

To identify VGW contigs that are still extending, contigs derived from iteration 30 were aligned against contigs from iteration 31 using BLASTn. If the top hit in iteration 31 was longer, then we determined which end of the original contig had extended based on the BLASTn HSP. We examined coverage by extracting data from the samtools depth file in a 60bp window from the end of each contig. The input cDNA transcripts were mapped against the contigs using GMAP, only the exon positions used to derive that VGW scaffold were used. Since the VGW scaffolds had been previously merged, there were some contigs with no exons mapped, these were excluded from the length analyses. These data were combined to analyse which contig ends had stopped extending, their median coverage depth and the length walked from the nearest mapped exon.

### Repeat identification and masking

The merged scaffolds from VGW, assembled chromosome 13 and 14 contigs and the BACs were processed through RepeatModeler to generate a classified consensus library of 2,103 sequences (Smit et al. 2014). This library was used in conjunction with RepeatMasker to mask repeats in our VGW scaffolds and across the BACs (Tarailo-Graovac and Chen 2009). The length of each repeat was determined from the GFF file, irrespective of potential overlaps.

### BAC comparison

To assess the quality of VGW assemblies we aligned the scaffolds and cDNA sequences against the 24 known BACs using BWA (Table S1) (Li 2013). Within this BAC collection there contains an apparent recent gene duplication; EU686400 and EU686411 are 99% identical over 62kb. Only one transcript in our collection corresponds to these two genes, and it is more similar to the EU686400 annotated transcript. We therefore excluded EU686411 from further analysis. Five of the remaining 23 BACs had no annotated transcript, although we were able to find VGW scaffolds for three of these (Table S1). Seven BACs had no corresponding transcript within our annotated CDS collection.

Percent identity values were calculated directly from the BAM files and contiguity was determined from mummer dotplots (Delcher et al. 2002).

**Table S1.** The BAC results. *EU686411 is a recent duplication of EU686400. **The fragmented BAC is incorrectly assembled/ordered.

| BAC | Gaps in BAC? | Annotated | Our cDNA | Contiguous ? | Exons (on BAC) | Mean percent identity |
| --- | --- | --- | --- | --- | --- | --- |
| JF490009 | N | thrsp | - | - | - | - |
| JF490010 | Y | nisch | ax_524095 | Y | 21 (1) | 98.4 |
| JF490011 | Y | - | ax_528389 | N** | 16 (7) | 97.1 |
| JF490012 | Y | arfip1 | ax_540425 | Y | 10 (1) | 98.9 |
| JF490013 | Y | adpgk | ax_1050 | Y | 7 (6) | 97.7 |
| JF490014 | Y | col1a2 | ax_526750 | Y | 52 (52) | 99.2 |
| JF490015 | Y | c-myc | ax_541715 | Y** | 3 (3) | 99.6 |
| JF490016 | N | nanog | ax_552852 | Y | 4 (4) | 100.0 |
| EU686400 | N | CALR1 | ax_4938 | Y | 9 (9) | 99.7 |
| EU686401 | N | NUDT1 | ax_542364 | Y | 7 (6) | 98.2 |
| EU686402 | N | AxNovel2 | - | - | - | - |
| EU686403 | N | - | - | - | - | - |
| EU686404 | N | HMGCR | ax_528625 | Y | 20 (19) | 99.2 |
| EU686405 | N | AxNovel3 | - | - | - | - |
| EU686406 | N | - | ax_558334 | Y | 2 (2) | 97.0 |
| EU686407 | N | plastin | ax_3218 | Y | 15 (2) | 99.5 |
| EU686408 | N | P2RX3 | - | - | - | - |
| EU686409 | N | - | ax_560001 | Y | 1 (1) | 99.3 |
| EU686410 | Y | RARRES | ax_548715 | N** | 6 (3) | 97.9 |
| EU686411 | Y | CALR | ax_4938* | - | - | - |
| EU686412 | Y | MIG-6 | ax_565889 | Y | 3 (3) | 99.8 |
| EU686413 | Y | Enolase 1 | ax_34716 | Y** | 4 (1) | 99.6 |
| EU686414 | Y | TIMP4L | - | - | - | - |
| EU686415 | Y | - | - | - | - | - |

### Human comparison and synteny

All chromosomal protein-coding transcripts from Ensembl release 89 (May 2017) were downloaded and made into a BLAST database of 80,434 cDNAs (Cunningham et al. 2015; Altschul et al. 1990). This database was searched against to find the human ortholog using a BLASTn e-value of 1e-10. For axolotl transcripts which identified the same human ortholog, we selected the axolotl transcript with the longest mean intron length. Information on human introns, exons and chromosomal location were extracted using the Ensembl API. Exon boundary sites were compared using a multiple alignment (muscle) of corresponding transcript sequences (Edgar 2004). The same protocol was used to compare human against axolotl, as to compare human and *Xenopus tropicalis.* To ensure that we were comparing like with like, exon boundaries that were not present in the opposing species were excluded as well as genes with a mean distance greater than 50 (939 axolotl genes and 232 *Xenopus tropicalis* genes) Large-scale synteny between human and chicken was obtained directly from the Ensembl website.

## Data Access

The VGW scaffolds, input transcripts and annotation files are available to download at figshare (https://tinyurl.com/y8gydc6n). The VGW program and accompanying python scripts are available at github (https://github.com/LooseLab/iterassemble) and include running instructions and an example dataset.

## Acknowledgements

The authors wish to thank Garry Morgan for helpful discussions and all those that contribute to open source software without whom this work would not be possible. Particular thanks are extended to Heng Li and Pierre Lindenbaum for their tools and benchmarking regarding extraction of fastq reads. This work was funded by the MRC, grant code MR/N020979/1.

### Disclosure Declaration

We have nothing to disclose.

## Supplementary Figures

**Figure S1.**
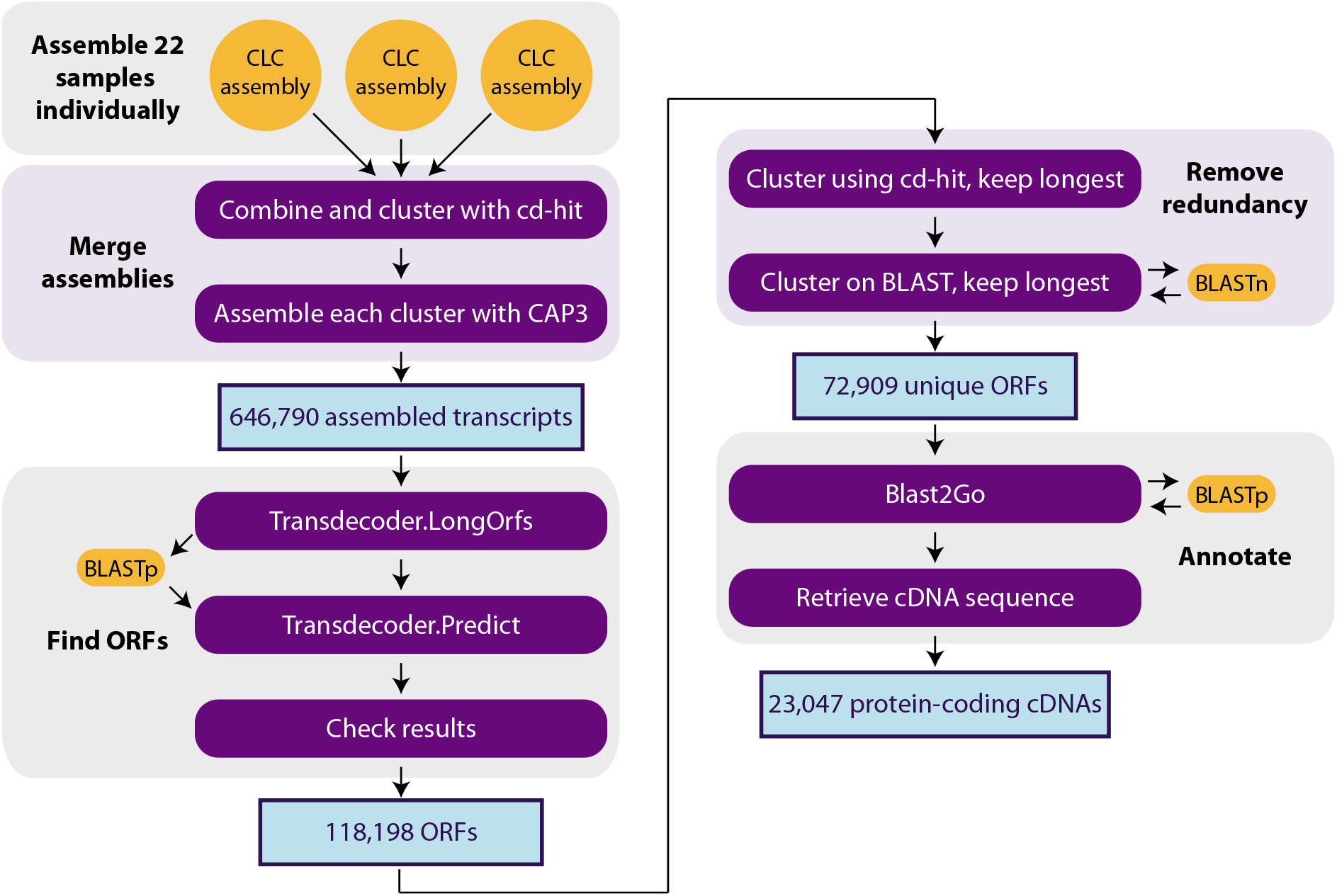
Transcriptome assembly pipeline. 22 RNA-seq samples of axolotl oocytes and early embryos were assembled following this protocol to form a final collection of 23,047 unique protein-coding cDNAs.

**Figure S2.**
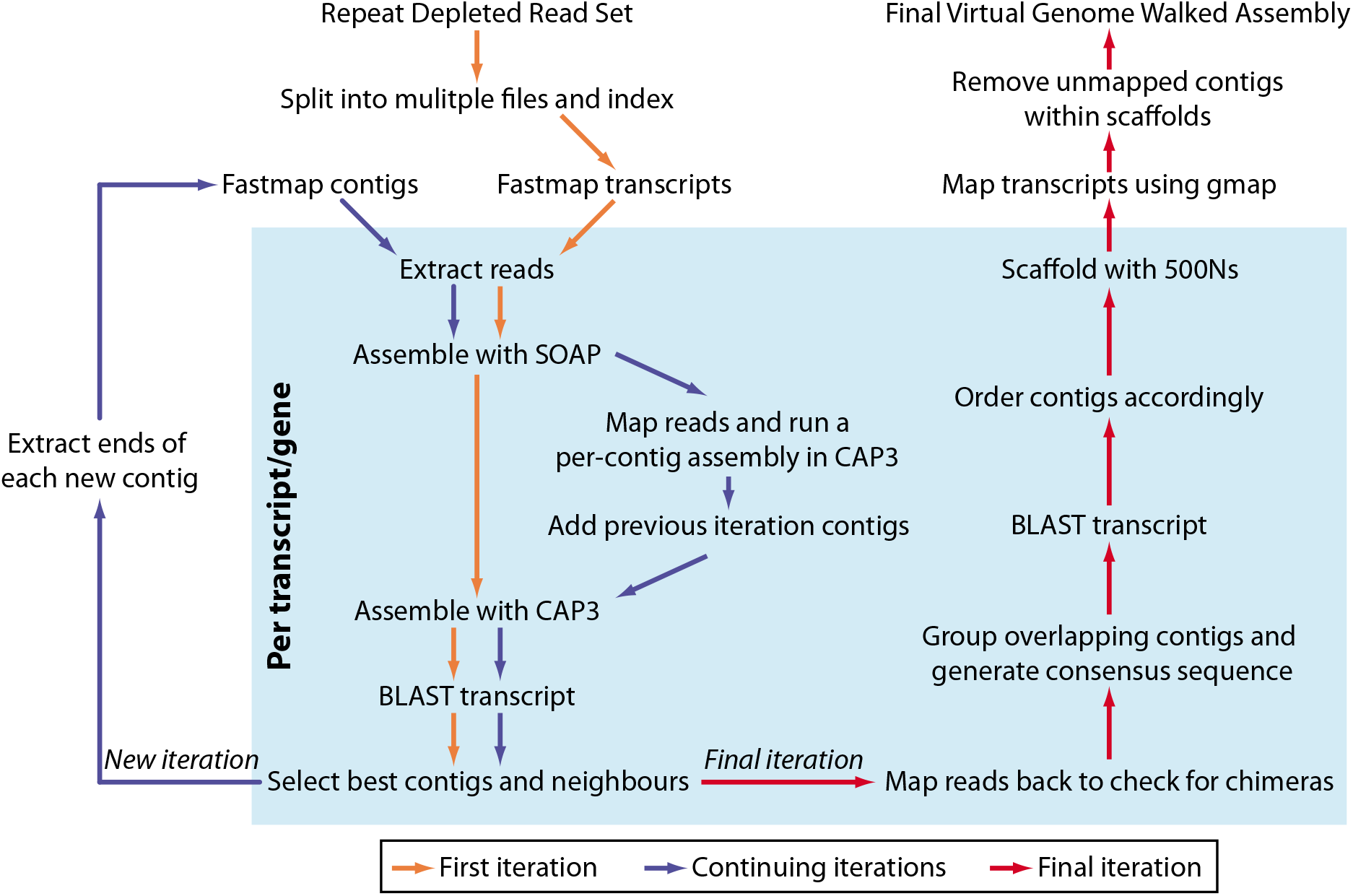
Detailed workflow of the Virtual Genome Walking pipeline. The process was altered depending on which iteration was ongoing (colored arrows). The steps in the blue box were run in parallel on each transcript/gene.

**Figure S3.**
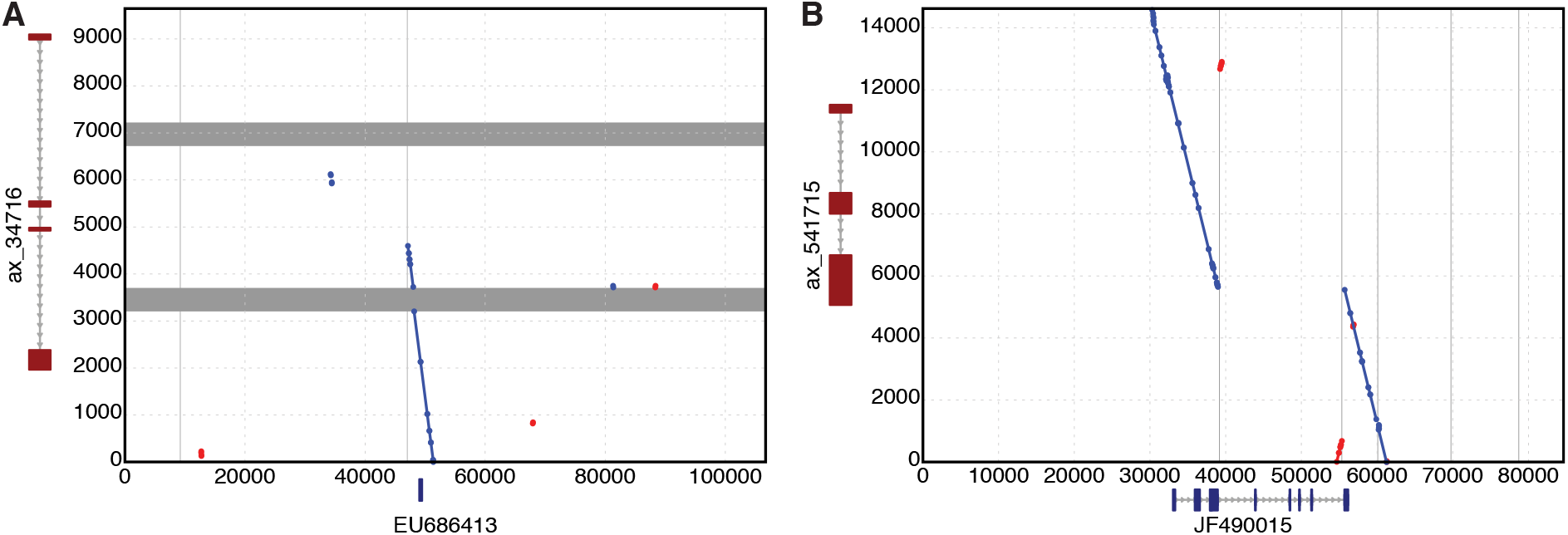
Dot-plots of the two VGW-BAC comparisons summarised in Figure 4C. Grey bars represent unassembled fragments and are shown to scale. The GMAP identified exons are shown alongside each input scaffold.

**Figure S4.**
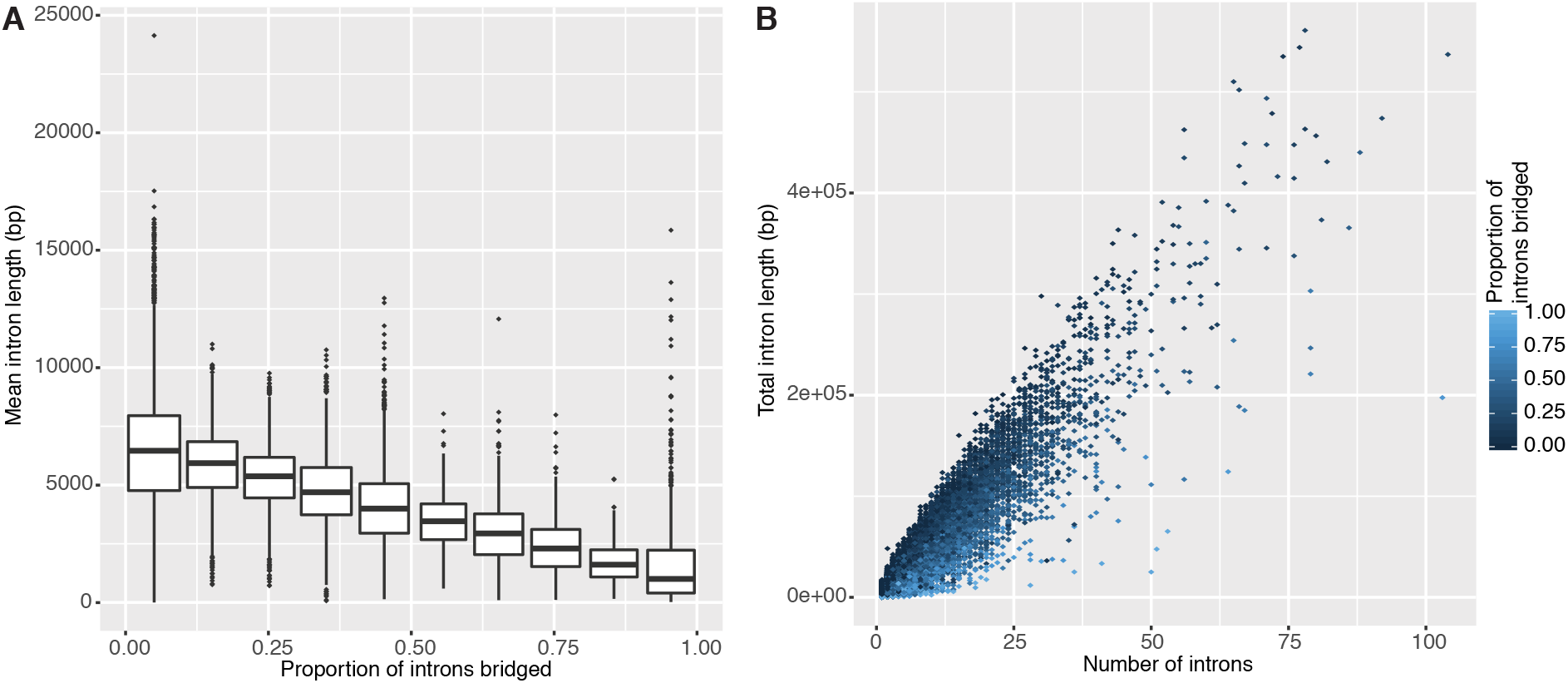
Axolotl intron lengths. A) Boxplots of mean intron length vs. proportion of introns able to be bridged. B) Total intron length correlates with the number of introns, the genes VGW can bridge tend to have a short total length and fewer introns.

**Figure S5.**
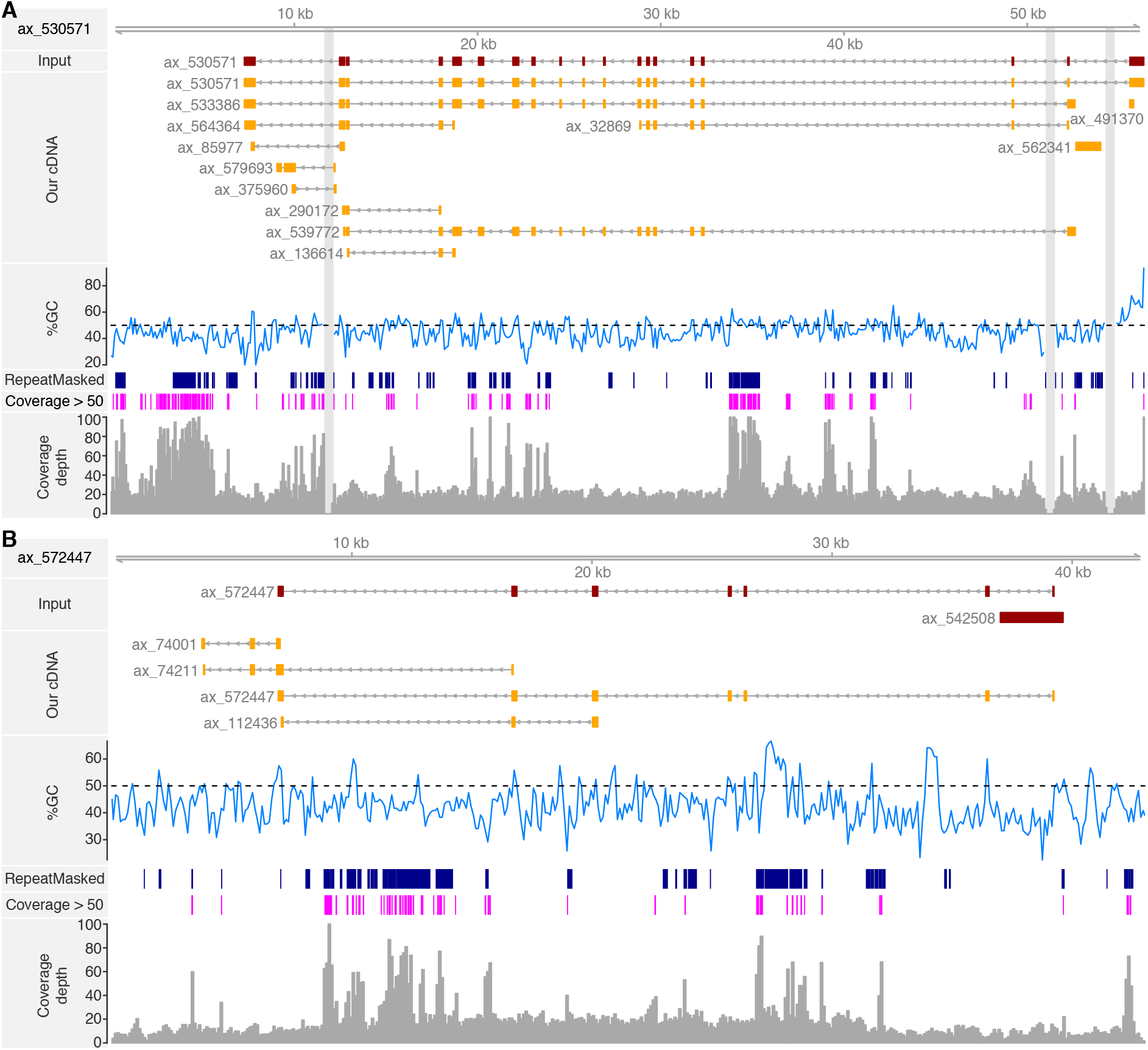
Examples of assembled axolotl introns. (A) The VGW output for ax_530571 with a 16Kb bridged intron and (B) ax_572447 a single contig of over 40Kb.

**Figure S6.**
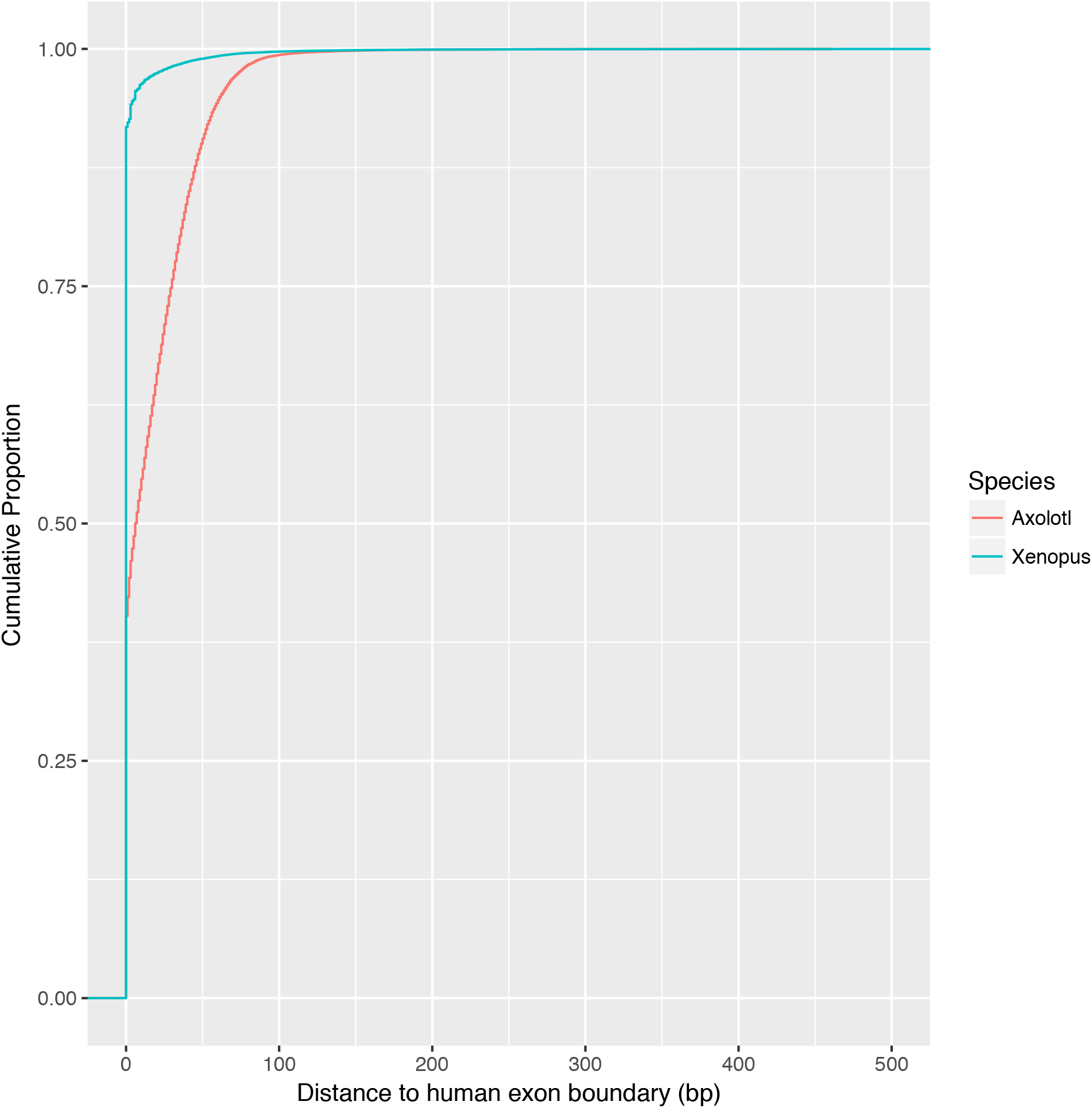
The exon-exon boundaries for orthologous transcripts compared to human for axolotl (red) and Xenopus (green).

**Figure S7.**
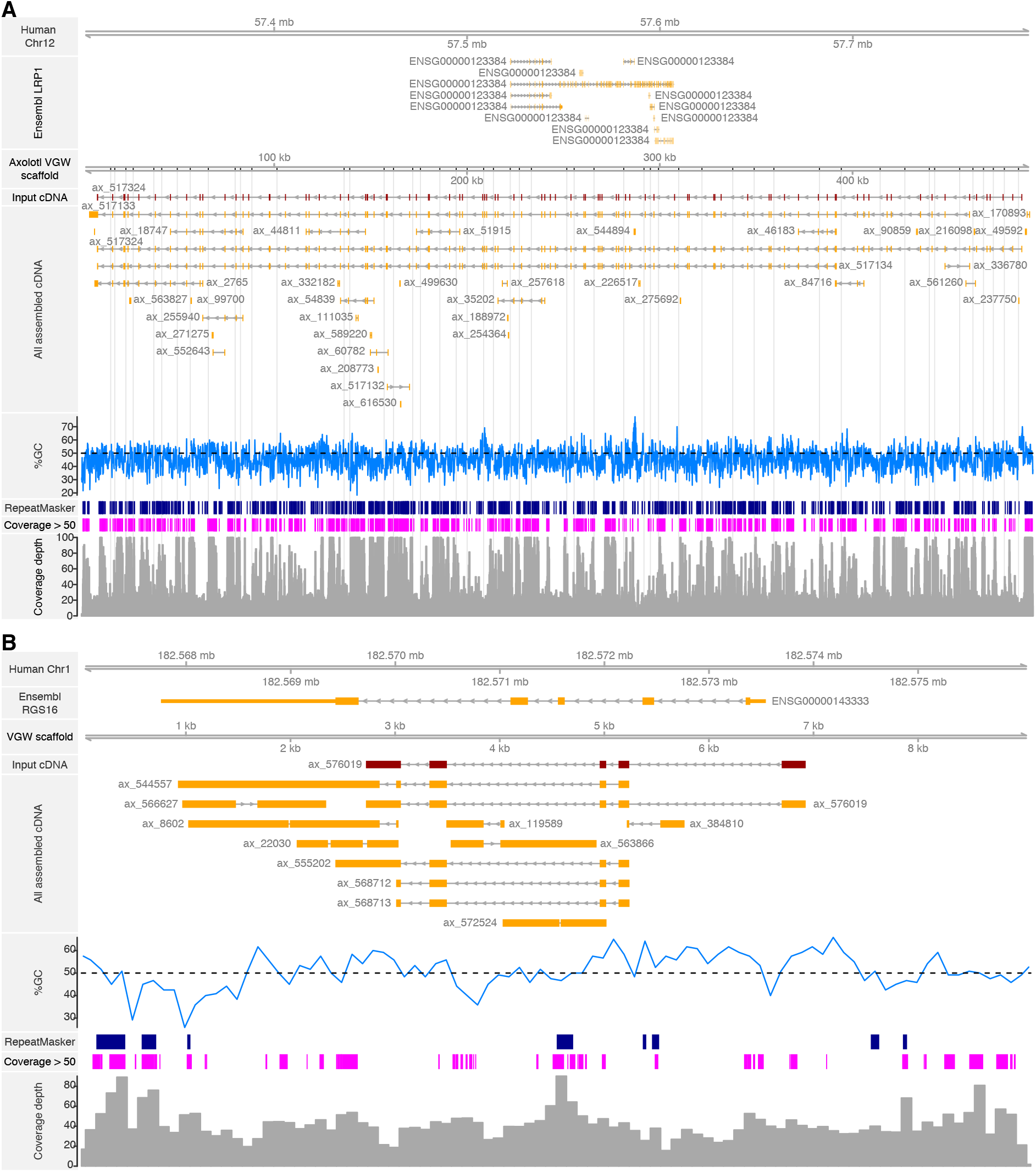
The VGW output for (A) ax_517324 and (B) ax_576019 compared to their human orthologs. (A) The axolotl gene is considerably larger than its human ortholog, the second example (B) shows orthologs that are similarly sized. The human and axolotl genome fragments are shown on the same scale, grey bars represent unassembled contig breaks.

**Figure S8.**
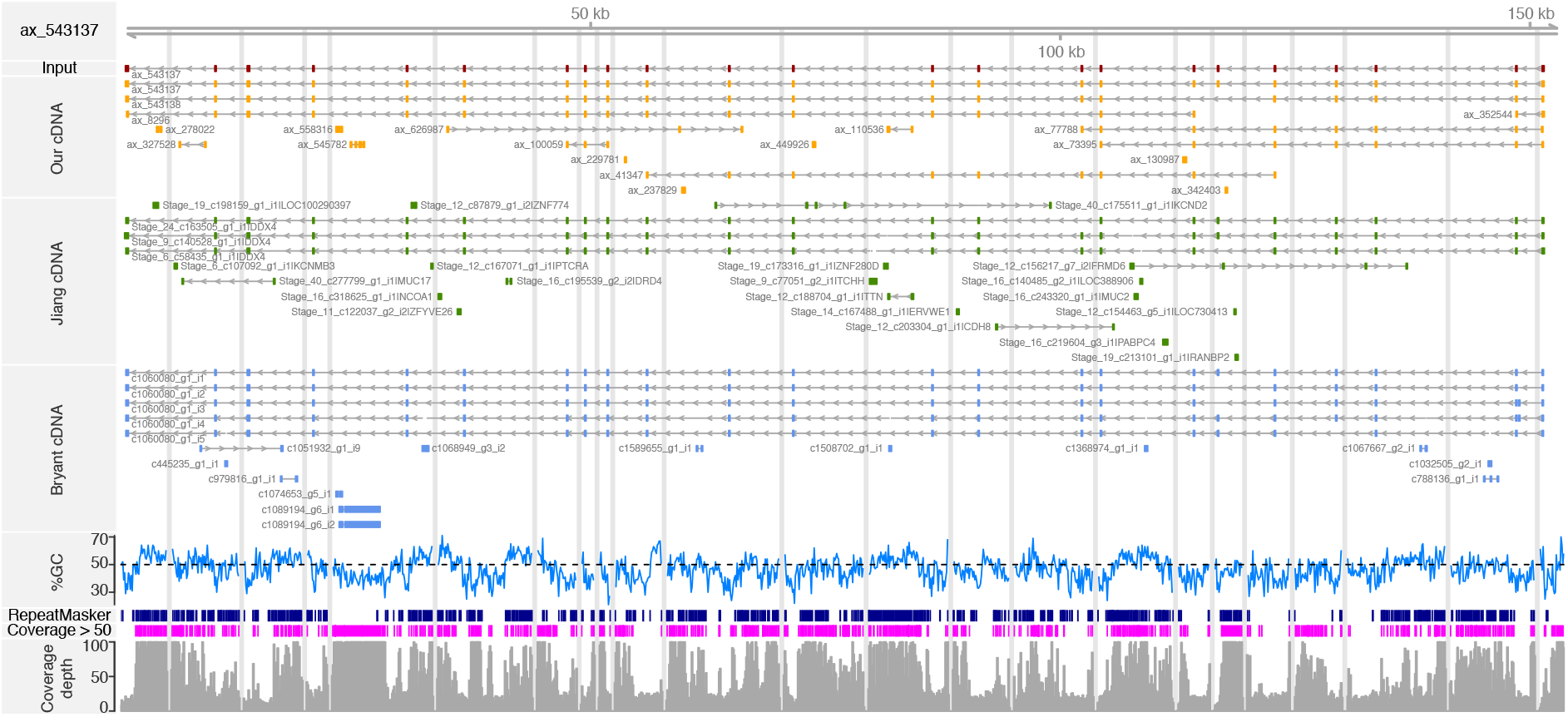
The VGW scaffold for DDX4 (ax_543137) is shown, vertical grey bars represent contig breaks. Transcripts assembled from our own data, alongside those from the Jiang and Bryant datasets are shown mapped to the scaffold (Jiang et al. 2017; Bryant et al. 2017). This demonstrates the multiple transcript variants of DDX4 expressed in axolotl. For visual simplicity, only those transcripts with at least two exons mapped are shown.

**Figure S9.**
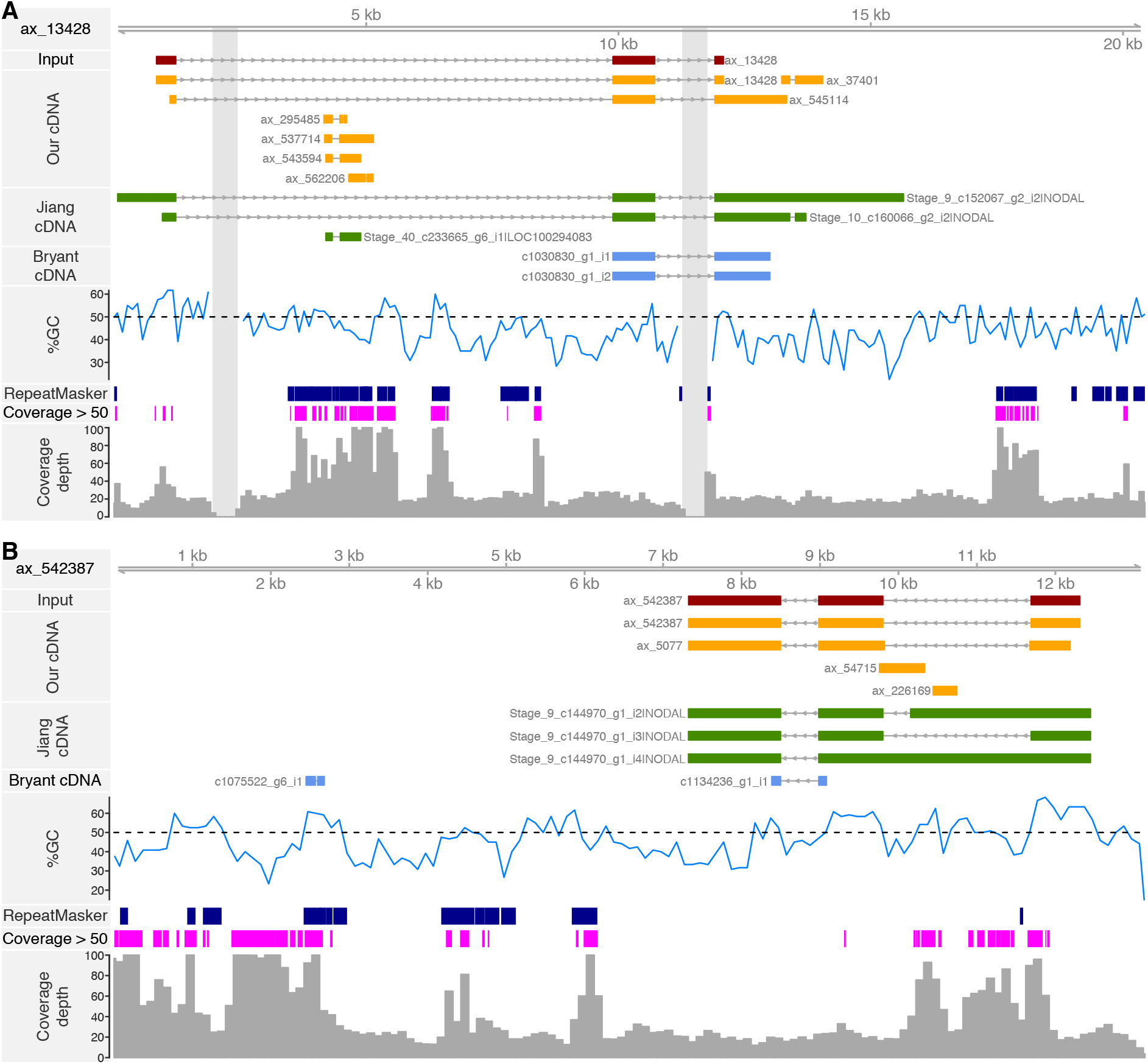
The VGW output for both NODAL genes. The two transcripts have assembled separate genome scaffolds and show two different gene models. Both are labeled as ‘NODAL’ in the Jiang dataset as they were annotated according to the single human gene (Jiang et al. 2017). Neither gene was completely assembled in the Bryant dataset (Bryant et al. 2017).

**Figure S10.**
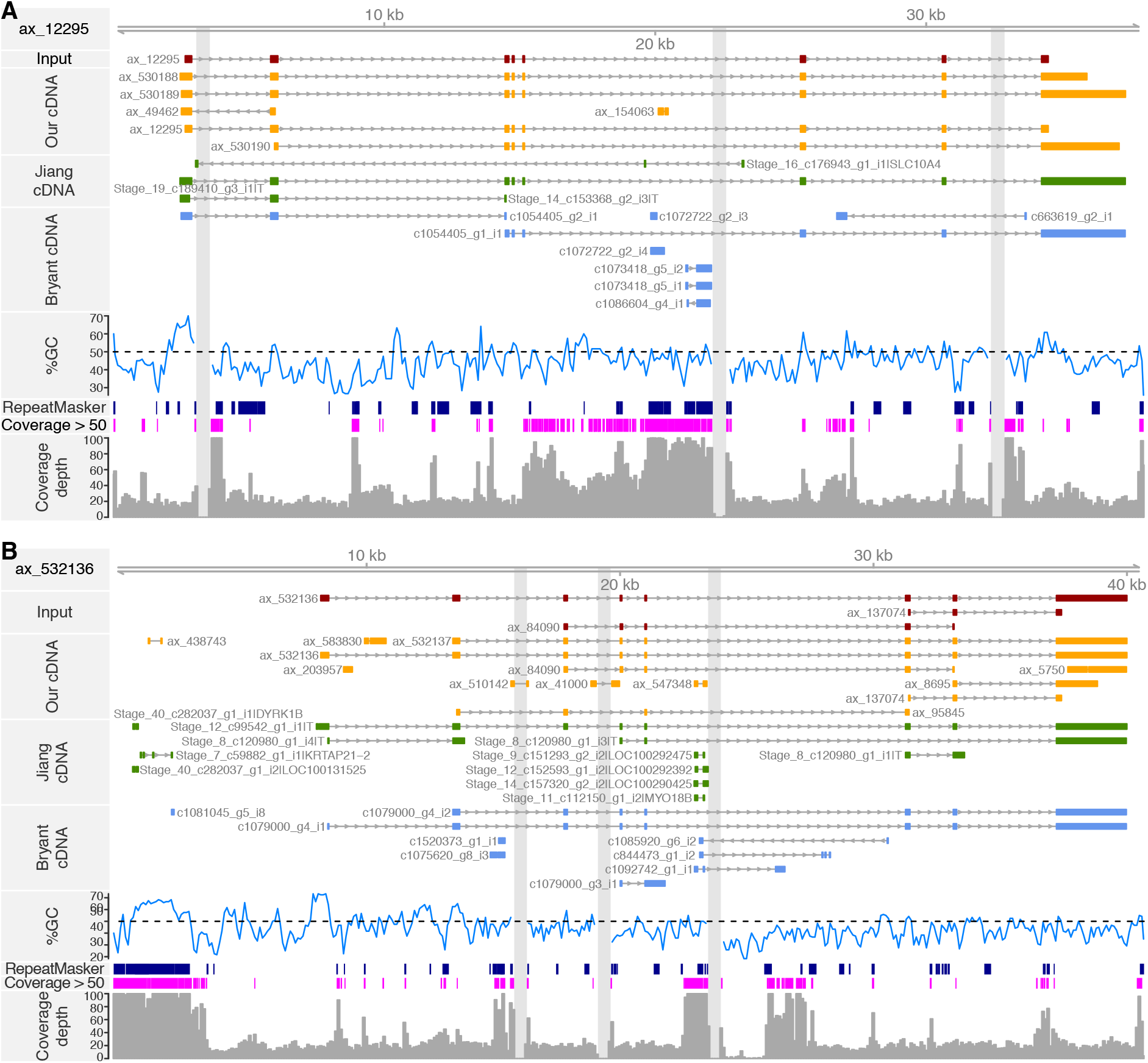
The VGW output for both Brachyury genes. These two transcripts have assembled different scaffolds and show two different gene models. Both are labelled as ‘T’ in the Jiang dataset as they were annotated according to the single human gene (Jiang et al. 2017).

**Figure S11.**
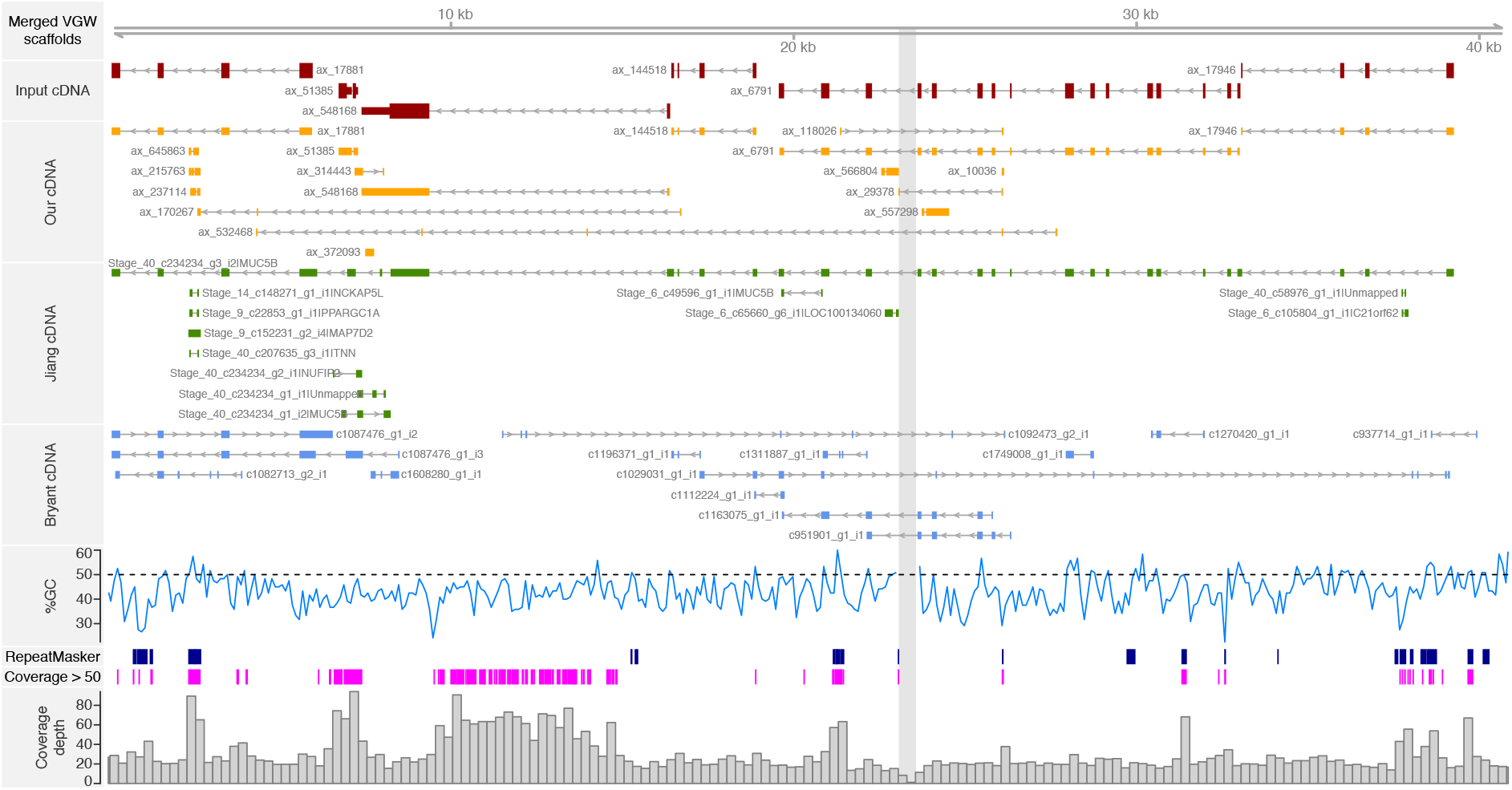
Three VGW scaffolds were merged, demonstrating the fragmented transcripts in our collection that are all derived from a single gene. A completely assembled transcript annotated as MUC5B was found in the Jiang dataset (Jiang et al. 2017).

**Figure S12.**
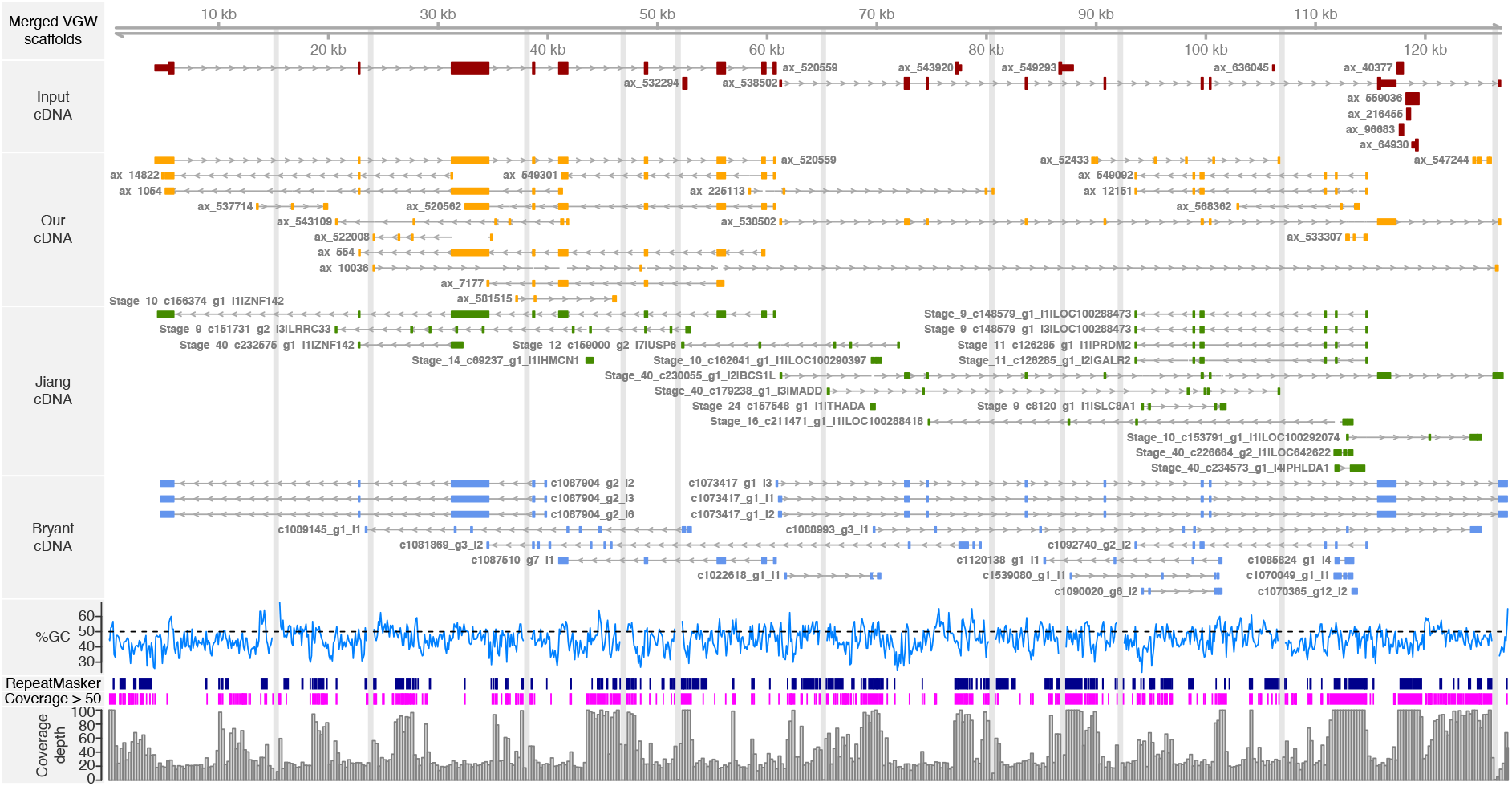
VGW was able to walk between these syntenic genes, ZNF142 and BCS1L. For clarity, only transcripts with at least four exons are shown in the all cDNA, Jiang and Bryant tracks (Jiang et al. 2017; Bryant et al. 2017).

**Figure S13.**
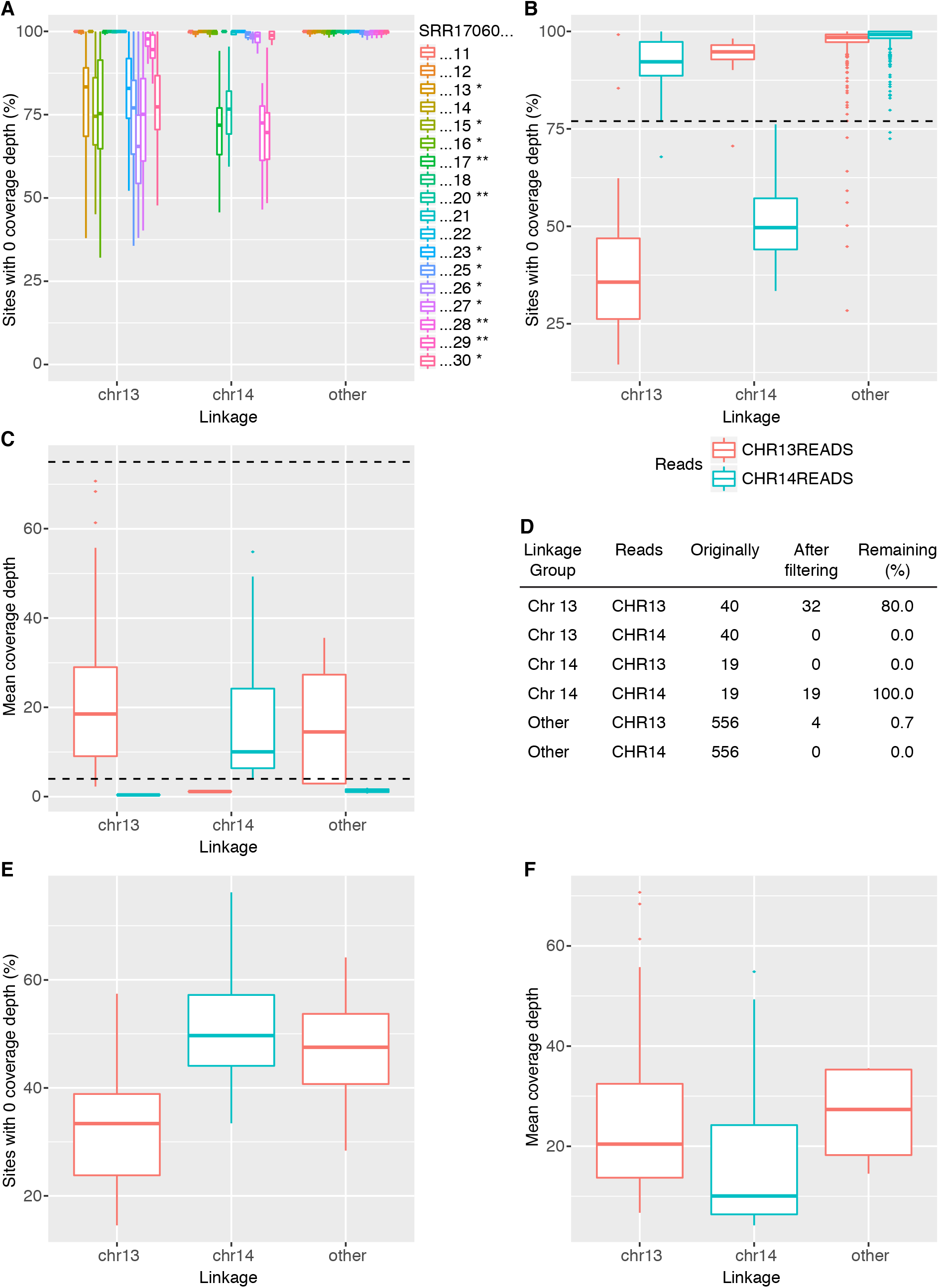
Filtering the Chr13 and Chr14 reads. A)The SRR libraries mapped to the linked exons, outliers are not shown for clarity. *Libraries assigned to AM13; ** libraries assigned to AM14. B) To distinguish AM13 and AM14 from other genes we removed those with more than 77% of exon sites having a coverage of 0 (dashed line). C) We also removed those with a mean depth of coverage less than 4 or greater than 75 (dashed lines). D) This removed a large proportion of genes linked to other chromosomes, and retained ~90% of the desired genes. We were unable to cleanly distinguish AM13 and AM14 from genes on other linkage groups based on sites with a coverage depth of 0 (E), or mean coverage depth (F).

**Figure S14.**
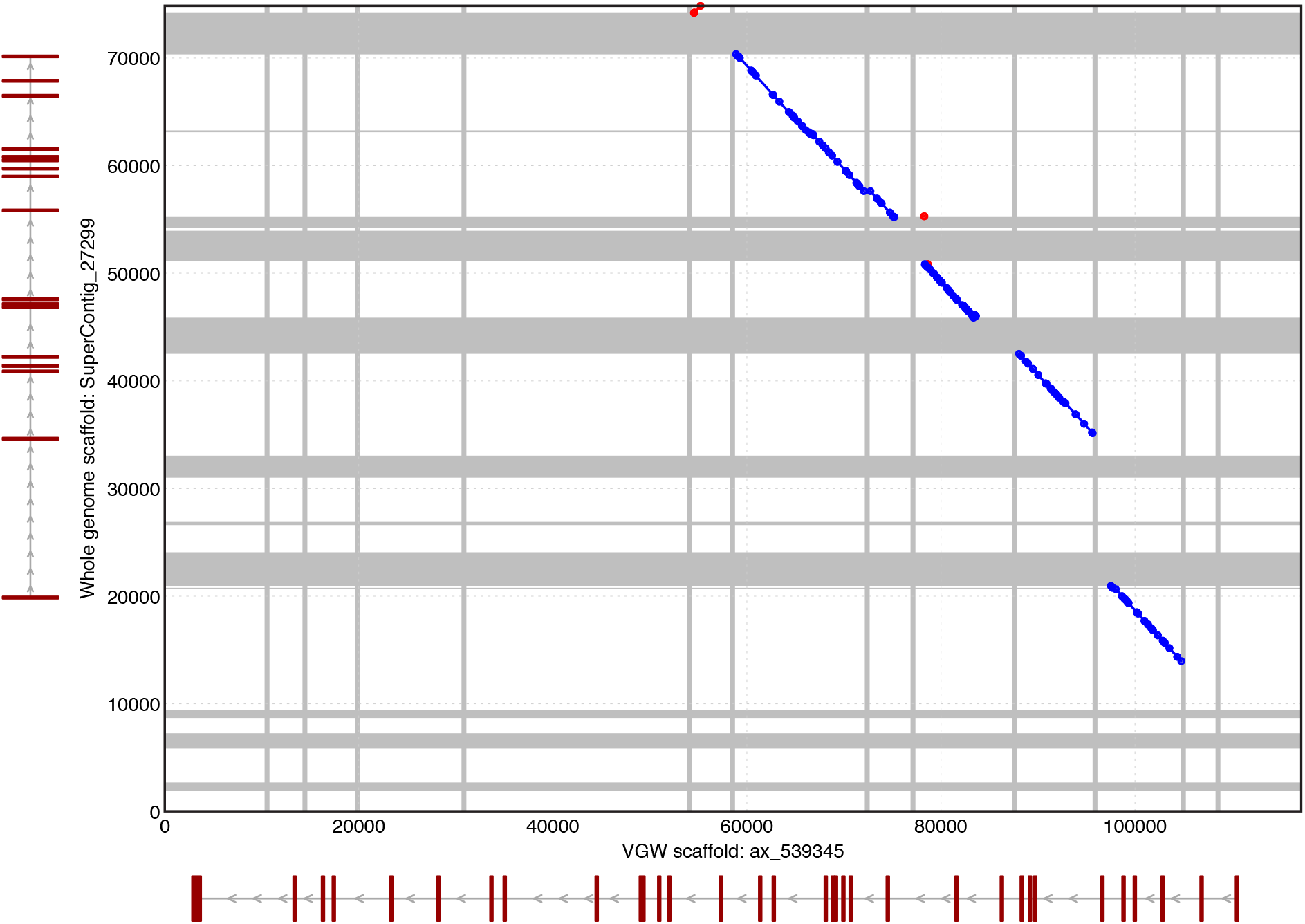
Dotplot comparison between one of our VGW scaffolds and one of the whole genome scaffolds. Regions with at least 100 Ns are shown in grey. The exon positions as calculated by GMAP are shown against each scaffold in red.

